# CD44 and Ezrin restrict EGF receptor mobility to generate a novel spatial arrangement of cytoskeletal signaling modules driving bleb-based migration

**DOI:** 10.1101/2024.12.31.630838

**Authors:** Ankita Jha, Ankit Chandra, Payam Farahani, Jared Toettcher, Jason M. Haugh, Clare M. Waterman

**Author notes:** Correspondence to, National Heart, Lung and Blood Institute National Institutes of Health, Building 50 South Drive Room 4537 MSC 8019, Bethesda, Maryland 20892-8019 United States.

## Abstract

Cells under high confinement form highly polarized hydrostatic pressure-driven, stable leader blebs that enable efficient migration in low adhesion, environments. Here we investigated the basis of the polarized bleb morphology of metastatic melanoma cells migrating in non-adhesive confinement. Using high-resolution time-lapse imaging and specific molecular perturbations, we found that EGF signaling via PI3K stabilizes and maintains a polarized leader bleb. Protein activity biosensors revealed a unique EGFR/PI3K activity gradient decreasing from rear-to-front, promoting PIP3 and Rac1-GTP accumulation at the bleb rear, with its antagonists PIP2 and RhoA-GTP concentrated at the bleb tip, opposite to the front-to-rear organization of these signaling modules in integrin-mediated mesenchymal migration. Optogenetic experiments showed that disrupting this gradient caused bleb retraction, underscoring the role of this signaling gradient in bleb stability. Mathematical modeling and experiments identified a mechanism where, as the bleb initiates, CD44 and ERM proteins restrict EGFR mobility in a membrane-apposed cortical actin meshwork in the bleb rear, establishing a rear-to-front EGFR-PI3K-Rac activity gradient. Thus, our study reveals the biophysical and molecular underpinnings of cell polarity in bleb-based migration of metastatic cells in non-adhesive confinement, and underscores how alternative spatial arrangements of migration signaling modules can mediate different migration modes according to the local microenvironment.

## Introduction

Cancer cell migration plays a crucial role in invasion and metastasis, which is often fatal^1–3^. During metastasis, tumor cells migrate through their surrounding microenvironment, navigating various tissue and extracellular matrix (ECM) geometries, including confined spaces, to spread to distant sites^4,5^. For cells to undergo motility, they must establish spatial polarity to create an actomyosin architecture that generates protrusion in the direction of migration, adhesion to the microenvironment, traction forces to move the cell relative to the environment, and retraction of the cell rear to make forward progress^6–8^. Tumor cells can employ several distinct modes of migration, which are influenced by their interactions with their particular microenvironment^9^. Depending on the availability of adhesion ligands, cancer cells can dynamically switch between integrin-mediated mesenchymal migration and adhesion-independent amoeboid migration^9–12^. In integrin-dependent migration, ECM ligand binding induces signaling that drives actin polymerization-based protrusions including lamellipodia and filopodia, and utilizes integrin-ligand binding and integrin linkages to the cytoskeleton to drive adhesion and traction generation^13^. However, when ECM ligand is unavailable or integrin signaling is suppressed, metastatic tumor cells fail to make actin-driven protrusions and instead switch to ameboid migration in which they adopt a bleb-driven mode of protrusion^14–16^. In bleb-based migration, hydrostatic pressure causes a directional bubbling out of the plasma membrane from the actin cortex in the direction of migration, and non-specific friction with the confined microenvironment together with actomyosin-driven cortical flow of transmembrane proteins drives adhesion and traction^17–20^. Blebbing is associated with increased Rho activity and is particularly pronounced in confined spaces, where high Rho-mediated contractility and elevated hydrostatic pressure from the confinement lead to the formation of large blebs that facilitate persistent migration^14,17–22^. Despite these observations, the precise molecular mechanisms underlying bleb-based migration in confined environments remain unclear.

While it is reasonably well understood how cell polarity is established during mesenchymal migration^6,7,23^, how this is achieved during bleb-based ameboid migration is largely unknown. Establishment of polarity requires symmetry breaking, which can occur in response to asymmetric cues, including diffusible molecules like during chemotaxis, substrate-bound signals as in haptotaxis, and physical cues as in mechanotaxis^13,23,24^. In the absence of such directional cues, cells may spontaneously break symmetry due to cytoskeletal instabilities^25^. However, such cytoskeletal asymmetry must be reinforced and maintained by polarized signaling within the cell. For instance, in adhesion-based mesenchymal migration, integrin engagement with an ECM gradient activates local Rac1 signaling^26,27^ that generates branched actin-driven lamellipodial protrusions^28,29^ that have primed integrin receptors at their tips ^30^, thus reinforcing protrusion and adhesion in a specific direction. In addition, in mesenchymal chemotaxis, growth factor gradients result in receptor tyrosine kinase (RTK) and G protein-coupled receptor (GPCR) activation at the cell front^31^. This initiates a cascade of locally self-amplifying signaling events through PI3K, resulting in accumulation of phosphatidylinositol (3,4,5)-trisphosphate (PIP3) and Rac or Cdc42 activation to promote formation of branched or bundled actin protrusions at the cell front ^7^. Simultaneously, phosphatidylinositol (4,5)-bisphosphate (PIP2) and Rho GTPase activity at the cell rear drive formin-based actin bundle formation and myosin II activation, generating traction and rear retraction^7,32^. In contrast, cells migrating via bleb formation in high confinement undergo spontaneous symmetry breaking ^19,22,24,25^, where loose formin-based actin bundles form just proximal to the bleb tip ^16,33^ and dense actin networks, along with myosin II activation, dominate the rear of the bleb ^19,21,33^. This asymmetry in polymerization and contraction drives cortical actomyosin flow within the bleb, which when coupled to transmembrane proteins that generate friction on the tightly confined microenvironment, propels the cell forward^19,22,33^. However, the role of polarized receptor signaling in reinforcing polarity in the absence of integrin engagement in bleb-based migration remains unknown^16^.

One system for maintaining polarized cellular asymmetries is by spatial regulation of receptor signaling on the plasma membrane. Establishment and maintenance of spatially constrained domains of transmembrane receptors in the fluid plasma membrane can occur through multiple mechanisms. One of these mechanisms is by advective movement along with membrane or cortical flows driven by membrane tension gradients or by direct or indirect binding to flowing cortical actomyosin^22,34–36^, as occurs during establishment of body-axis polarity in the early C. elegans embryo^37^. Alternatively, regions of specific plasma membrane composition can be formed by local addition or removal of surface proteins by spatial regulation of endo- and exocytosis, as occurs in maintenance of apical and basal-specific ion channels and transporters in polarized kidney epithelia^38^ or like in *Drosophila melanogaster* ovary, where RTK signaling is spatially restricted to distinct regions of migrating border cells^39,40^. Finally, local surface receptor compartmentalization can also be mediated by polarizing endoplasmic reticulum-plasma membrane contact^41^ or spatial restriction of receptor lateral mobility within the plasma membrane, also known as “corralling”^42^. For instance, during phagocytosis, large transmembrane proteins, such as Cluster of Differentiation 44 (CD44), limit the diffusion of Fcγ receptors by corralling at the rear of polarized macrophages, leaving them more mobile at the leading edge^43^. Whether these mechanisms govern establishment of stable polarity in bleb-based migration is yet to be determined.

In this study, we aimed to identify the mechanisms that mediate the cell polarization that enables persistent bleb-based migration of metastatic cancer cells in non-adhesive confined microenvironments. Our previous work demonstrated that leader bleb-based migration in metastatic melanoma cells under confinement is driven by Erk signaling^20^, which is linked to EGF/EGFR/Ras/Raf pathways. We thus hypothesized that the polarized distribution of growth factor receptor tyrosine kinases and their downstream signaling components, such as phosphoinositides and Rho GTPases^44–46^, is required for maintaining persistent polarized migration in metastatic melanoma cells under confinement. To test this hypothesis, we utilized an *in vitro* cell confinement assay together with time-lapse high resolution imaging and optogenetic control of local signaling to dissect the mechanisms mediating the establishment and maintenance of cell polarity during bleb-based migration. We find that bleb-based migration occurs via spatial constraint of similar signaling modules utilized to drive polarity in lamellipodium-based chemotactic migration, only these modules are largely spatially inverted. Whereas in lamellipodial migration growth factor receptors and downstream PI3K-Rac signaling localize to the cell front, we find their localization to the bleb rear is required to maintain bleb stability and persistent migration. We further reveal that this rear-to-front EGFR gradient is established by restricting mobility of EGFR in the plasma membrane at the rear of the bleb by CD44 interacting with cortical actin via the linker protein ezrin. Our results thus identify a novel mechanism by which restricting EGFR-PI3K-Rac signaling axis to the back of the bleb results in rapid migration of cancer cells that may be critical to their highly invasive behavior in vivo.

## Results

### EGFR signaling through PI3K is required for leader bleb stability and persistent migration under non-adhesive confinement

Here we sought to understand the highly polarized bleb morphology of metastatic melanoma cells migrating in a confined, non-adhesive microenvironment. We first asked if this morphology was functionally relevant to cell migration. We performed time-lapse imaging of A375M2 human metastatic melanoma cells that were held in non-adhesive confinement to 3 μm height using a polydimethylsiloxane (PDMS) cellular confinement device, mimicking the degree of tissue confinement observed during metastasis *in vivo*^5^. We then characterized the relationship between cell morphology and motility. Melanoma cells confined in a non-adhesive environment exhibited blebs that were typified by distinct states of morphology and stability (Extended Figure 1A). This included a minor population (∼9 percent) of cells that exhibited small transient round blebs around their peripheries with a lifetime of less than a minute ((Figure 1A, top row, 1B)^17^. Confinement also induced elongated blebs (Fig 1A, rows 2-4), as described for many cell types from different organisms^19,20,22,47^.

**Figure 1:**
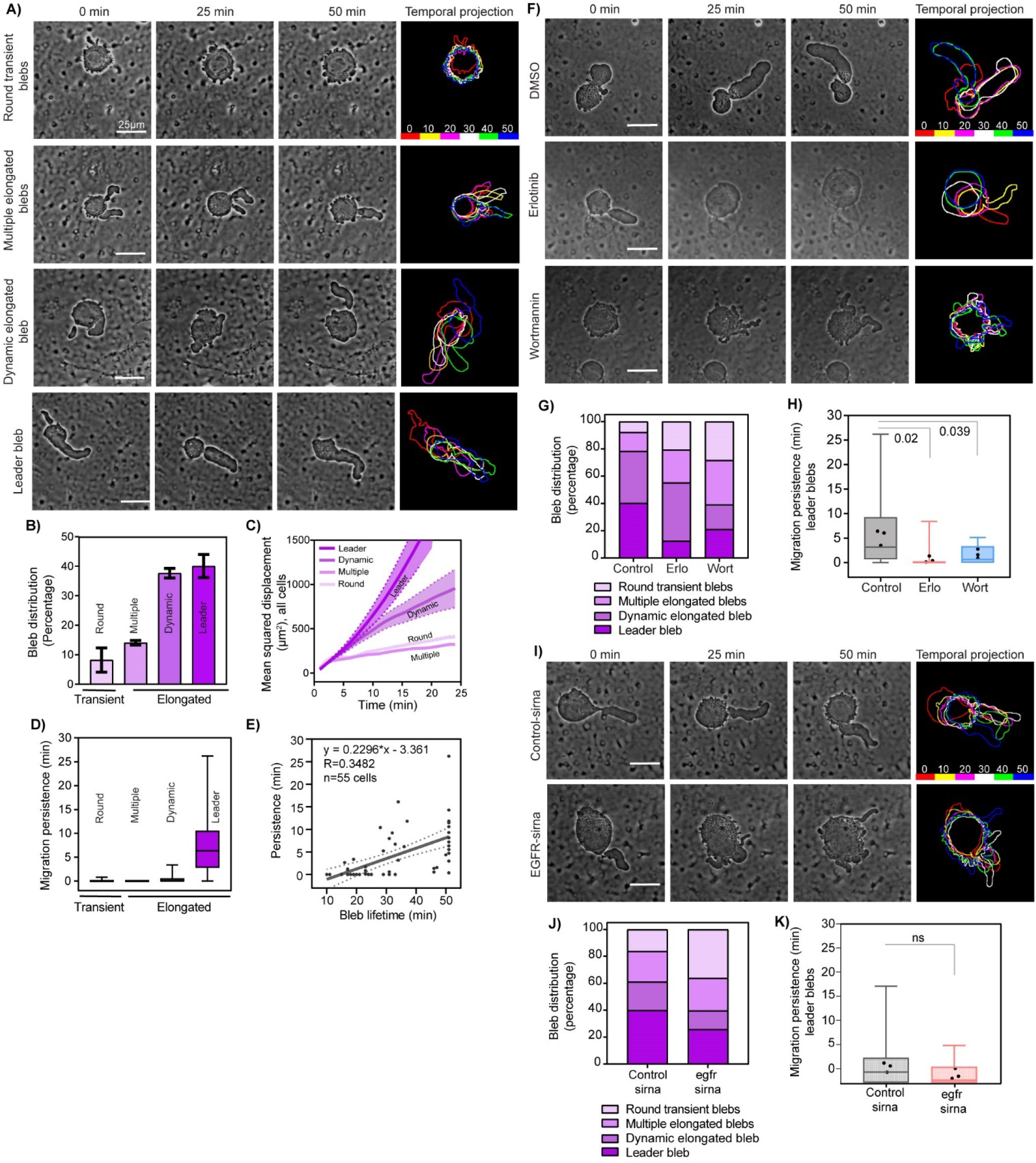
EGFR signals via PI3K to maintain leader bleb-based migration in non-adhesive confinement. A) Representative time-lapse phase-contrast images (left three columns) of A375M2 melanoma cells under 3 µm confinement. (A) Image showing four major bleb phenotypes: round transient blebs (top row), multiple elongated blebs (second row), dynamic elongated blebs (third row), and leader blebs (bottom row). Temporal color-encoded projection of cell shapes at 10-minute intervals (right column) of the cells shown at left. (B) Quantification of bleb phenotype distribution percentage. N= 360 cells from 3 experiments. (C) Round transient blebs (n= 21 cells), multiple elongated blebs (n= 18 cells), dynamic elongated blebs (n= 31 cells), and leader blebs (n= 44 cells) from N=3 experiments. Mean-square displacement over time with average (dark line) and standard deviation (dotted line, shaded area) shown (D) Box plots of migration persistence across different bleb phenotypes with round transient blebs (n=15 cells), multiple elongated blebs (n=17 cells), dynamic elongated blebs (n=25 cells), and leader blebs (n=29 cells) from N=3 experiments. (E) Migration persistence of cells with polarized elongated blebs versus bleb lifetime (n=55 elongated blebs). (F) Cells under confinement treated with DMSO, erlotinib (50nM) or wortmannin (10μM). (G) Quantification and stacked column bar graph of blebbing phenotypes after drug treatments as in (F) (DMSO n=360 cells; erlotinib n=208 cells and wortmannin n=166 cells from N=3 independent experiments). (H) Box plot of migration persistence of cells with leader blebs in drug treated cells as in (F), (DMSO n=37 cells; erlotinib n=23 cells and wortmannin n=19 cells from N=3 experiments). (I) Cells under confinement after non-targeting (Control-siRNA) or smart pool siRNAs targeting EGFR (EGFR-siRNA). (J) Quantification of blebbing phenotypes post-siRNA treatment (Control-siRNA n=118 cells, EGFR-siRNA n=116 cells) (K) Migration persistence of cells with leader blebs after control (n=42 cells) or EGFR-siRNA (n=22 cells) treatment. Box plots in (H, K) show the median value and box extending from the 25^th^ to 75^th^ percentiles and whiskers show the minimum and the maximum values of the data. P value, not significant (P<0.05, ns), Welch’s t-test.

Quantitative analysis showed that cells with elongated blebs could either be in a non-polarized state with multiple elongated blebs around their peripheries (∼13 percent) or in a highly polarized morphology with a single elongated bleb (∼75 percent) (Figure 1B). Cells with single elongated blebs exhibited two distinct types that we will refer to as dynamic elongated blebs (Extended Figure 1A), which were those that extended and retracted with a lifetime of <25 minutes (∼35 percent) and leader blebs, which were stable and had a lifetime of >25 min (∼40 percent of cells; Figure 1A, B; Movie 1)^20^. To determine if different bleb morphologies were functionally relevant to migration, we analyzed motility parameters for each category of cell shape in confinement. We found that cells exhibiting stable leader blebs migrated much more efficiently compared to cells exhibiting round transient blebs, multiple elongated blebs, or a dynamic elongated bleb, displaying significantly greater migratory range, as measured by a greater slope on a mean squared displacement over time plot (Figure 1C), migration persistence (Figure 1D) and higher diffusion coefficient (Extended Figure 1B). Indeed, plotting the lifetime of elongated blebs before their retraction vs either their migration persistence (Figure 1E) or diffusion coefficient (Extended Figure 1C) showed significantly positive correlations. In contrast, similar analysis of bleb lifetime vs speed showed that cells with an elongated bleb moved at similar speeds, independent of bleb lifetime (Extended Figure 1D and 1E). These results show that stabilization and maintenance of a polarized, elongated, leader bleb morphology is critical to efficient and persistent migration of metastatic melanoma cells in non-adhesive confinement.

We then sought to investigate the molecular basis underlying stabilization of highly polarized leader blebs in melanoma cells in non-adhesive confinement. As sensing of soluble growth factors in the microenvironment is critical to cell polarization in directional, adhesion-based migration of cancer cells^48,49^, we probed the role of serum components and growth factors in leader bleb polarization and stability. We observed that confinement of serum-starved A375M2 cells resulted in a strong decrease in formation of stable leader blebs and increased the fraction of cells with round transient blebs compared to cells confined in serum-containing media (Extended Figure 1F, G). Analysis of cell motility parameters showed that serum starvation strongly reduced migratory range (Extended Figure 1H) and the few cells that exhibited leader bleb morphology had reduced migration persistence, but no effect on cell speed (Extended Figure 1I, J). As EGF signaling is critical to cell migration and developing drug resistance in metastatic melanoma^50,51^ and starvation causes downregulation of this signaling (Extended Figure 1K), we reasoned that EGF could be the serum factor responsible for promoting leader bleb-based migration in non-adhesive confinement. Importantly, addition of EGF to serum-starved cells partially rescued the leader bleb morphology (Extended Figure 1F, bottom panel and Extended Figure 1G; Movie 2) and increased the migratory range and migration persistence (Extended Figure 1H, 1I) but did not impact the cell speed (Extended Figure 1J). We found that pharmacological inhibition of either EGFR tyrosine kinase activity using erlotinib (10μM) or inhibition of its downstream target phosphatidylinositol 3-kinase (Pi3K) with wortmannin (10nM) strongly decreased the proportion of cells with stable leader blebs (Figure 1F and 1G; Movie 3) and their migratory range (Extended Figure 1L). Furthermore, those cells that did make leader blebs under either treatment showed significantly reduced migration persistence (Figure 1H), without a change in migration speed (Extended figure 1M). To substantiate the notion that EGF signaling promoted persistent leader bleb-based migration in non-adhesive confinement, we examined the effects of knocking down the EGF receptor by siRNA (EGFR-KD, Extended Figure 1N). This showed that compared to non-targeting siRNAs (NT), EGFR-KD strongly inhibited the formation of both dynamic and stable elongated blebs (Figure 1I, 1J; Movie 4J) and significantly reduced cell migratory range (Extended Figure 1O). Though the EGFR-KD cells exhibiting leader bleb morphology had both reduced migration persistence (Figure 1K) and speed (Extended figure 1P), these did not rise to statistical significance. Together, these results show that EGF signaling through PI3K is required for stabilization and maintenance of a polarized, elongated leader bleb morphology to drive persistent migration and promote migratory range under non-adhesive confinement.

### EGFR, PI3K, and Rac1 activities exhibit rear-to-front gradients in leader blebs

We next focused on understanding how EGF signaling mediates stabilization of the polarized morphology of elongated blebs during migration of melanoma cells under non-adhesive confinement. We examined the localization of EGFR by expressing EGFR-GFP and super-resolution confocal imaging. This revealed that a strong gradient of EGFR-GFP was present along the plasma membrane of leader blebs with a very low concentration at the distal bleb tip that rose to a peak at about a quarter of the length from the bleb base, and remained elevated at the neck where the bleb connects to the cells body (Figure 2A,B). Quantification of the polarity index across the bleb confirmed this rear-to-front gradient, with the bleb base exhibiting an average ∼2.5 fold enrichment of EGFR-GFP compared to the bleb tip (Figure 2C). This surprising rear-to-front polarity of EGFR on leader blebs led us to ask whether the localization of activated EGFR differed from that of the total EGFR, i.e. if there was a local difference in sensitivity to EGFR activation within the bleb. To approach this question, we utilized a modular biosensor in which the C-terminus of EGFR is fused with a tyrosine activation motif from CD3, such that when EGFR signaling is high, this motif is phosphorylated, resulting in the selective recruitment of the SH2 domain of ZAP70 (ZtSH2) to the membrane ^52^. Live imaging of cells co-expressing fusion red-labeled EGFR-CD3 (EGFR-CD3-FR) and clover-tagged ZtSH2 (ZtSH2-CLV) showed that both constructs formed strong rear-to-front gradients very similar to that seen for EGFR-GFP, indicating that EFGR activity is concentrated at the base of the leader bleb (Figure 2D, 2E). Quantification of the ratio of ZtSH2-CLV to EGFR-CD3-FR showed that while there appeared to be a slight elevation in the fraction of activated receptor towards the bleb tip, the polarity index of the ratio was close to zero, indicating that there was no spatial bias in sensitivity of EGFR (Figure 2F, 2G). Taken together, these results show that EGFR localization and activity are highly polarized in a rear-to-front gradient in leader blebs during migration in non-adhesive confinement.

**Figure 2:**
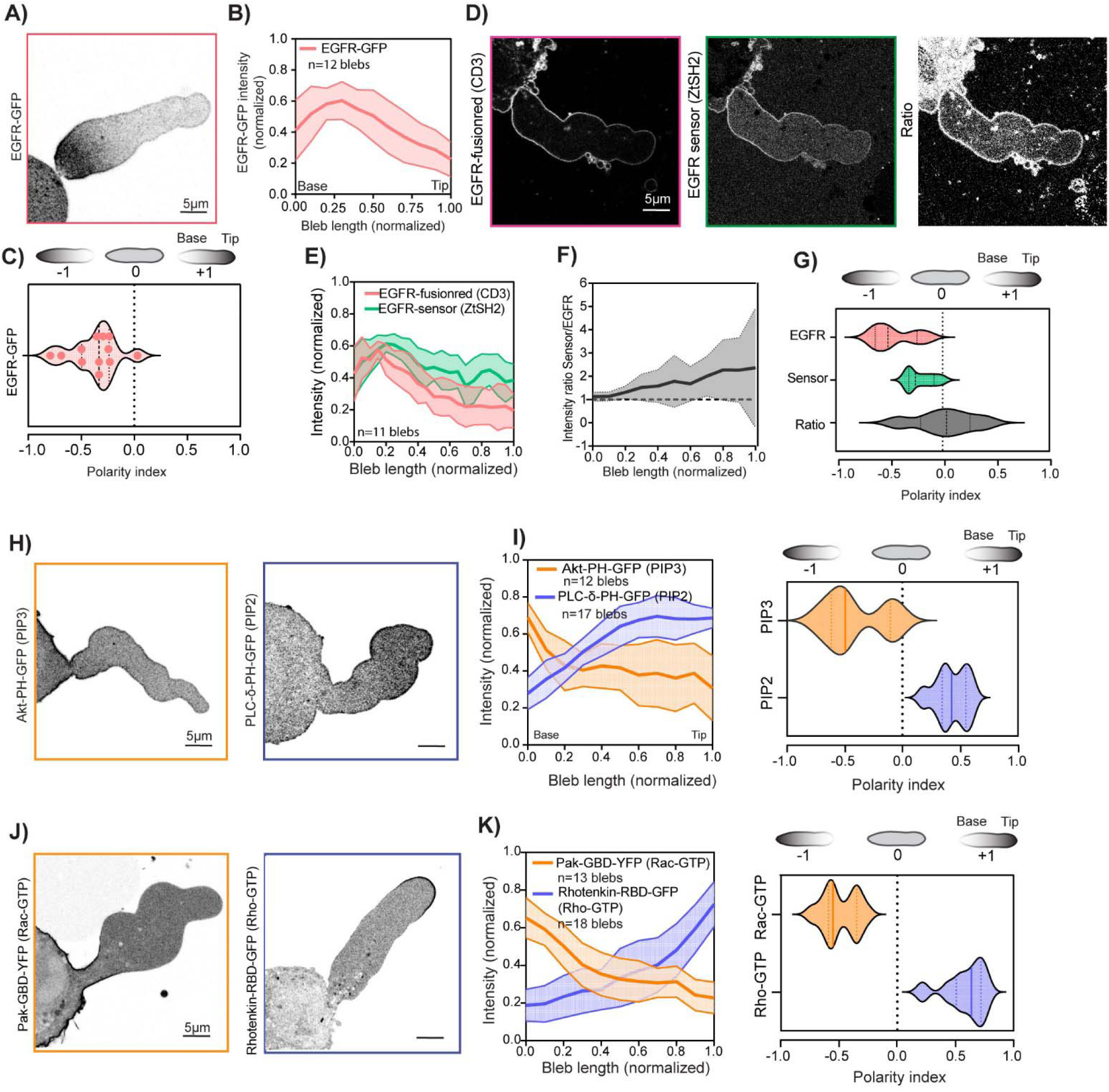
EGFR, PI3K and Rac1 activities form rear-to-front gradients in leader blebs. (A, D) Confocal images of representative leader blebs in A375M2 melanoma under 3μm confinement. (A) Inverted contrast image of a cell transfected with EGFR-GFP. (B,C) Graph showing the mean normalized intensity of EGFR-GFP (red box) across the normalized length of the leader bleb, mean ± s.d. (left) and violin plot of EGFR-GFP polarity index (right); n=12 cells. (D) Stable expression of EGFR-CD3-fusionred (red box, left) and ZtSH2-Clover (sensor, green box, center) and ratio image of sensor to EGFR-CD3-fusionred (right). (E,F) Quantification of mean normalized intensity of the constructs in (D) across the normalized length of leader blebs. (F) Normalized intensity ratio profile of EGFR to the sensor across the normalized length of the bleb (G) Violin plot of polarity indices of fluorescence distribution along the leader bleb of constructs from (D) and ratio, mean ± s.d., n= 11 cells. (H, I) Super resolution confocal images taken during confinement to 3 um of A375M2 melanoma cells. (H) Inverted contrast images of cells transfected with Akt-PH-GFP (orange box) and PLC-δ-PH-GFP (purple box). (I) Normalized intensity profile of Akt-PH-GFP (orange, n=12 blebs) or PLC-δ-PH-GFP (purple, n=17 blebs) across the normalized length of the bleb, mean ± s.d. (left) and violin plot of polarity indices of fluorescence distribution along the leader bleb of constructs from (H). (J) Inverted contrast images of cells transfected with Pak-GBD-YFP (orange box) and Rhotenkin-RBD-GFP (purple box). (K) Normalized intensity profile of Pak-GBD-YFP (orange, n=13 blebs) or Rhotenkin-RBD-GFP (purple, n=18 blebs) across normalized length of the bleb (left) and violin plot of polarity indices of fluorescence distribution along the leader bleb of constructs from (J) (right).

We next asked if signals downstream of EGFR also exhibit a polarized distribution during leader bleb-based migration. To examine the subcellular localization of PI3K signaling, we expressed either a fusion of the PH domain of Akt with GFP (Akt-PH-GFP) as a bio-sensor for phosphatidylinositol (3,4,5)-triphosphate (PIP3) localization ^53^ or a fusion of the PH domain of PLC-delta with GFP (GFP-C1-PLCdelta_PH) to report on phosphatidylinositol (4,5)-biphosphate (PIP2) ^54^, and imaged cells by confocal microscopy (Figure 2H).

Imaging and measurement of polarity index within leader blebs showed that, similar to EGFR and its activity proxy, Akt-PH-GFP (PIP3) exhibited a strong rear-to-front gradient with high intensity at the base of the bleb and low levels at the tip (Figure 2H, left panel, 2I). In contrast, PLC-delta-PH-GFP (PIP2) revealed a front-to-rear gradient the inverse of that of PIP3, with highest concentration at the tip of the leader bleb (Figure 2H, right panel, 2I). As PI3K signals to Rac1 ^27^ and PIP2 on the plasma membrane recruits RhoA and its regulatory proteins ^55^, we examined the localization of Rac1 and RhoA activities within leader blebs. We expressed the GTP binding domains (GBD) of either Pak fused to YFP (Pak-GBD-YFP) or Rhotekin fused to GFP (Rhotekin-RBD-GFP), which act as GTP-dependent binding sensors for Rac1 or RhoA, respectively. Confocal imaging showed that similar to EGFR-GFP and Akt-PH-GFP, Pak-GBD-YFP, and thus Rac1 activity, concentrated at the base of the bleb in a rear-to-front gradient (Figure 2J, left panel, 2K). In contrast, like PLC-delta-PH-GFP, Rhotekin-RBD-GFP, and thus RhoA activity, localized toward the bleb tip (Figure 2J right panel, 2K). Overall, these results show that cells undergoing migration in non-adhesive confinement spontaneously polarize EGFR in the plasma membrane of the leader bleb to generate strongly localized regions of subcellular signaling, with EGFR downstream effector PI3K promoting PIP3 accumulation and Rac1 activity in rear-to-front gradients, and their antagonists, PIP2 and RhoA activity, localized to the bleb tip. Furthermore, this rear-to-front polarization of signaling in leader blebs in non-adhesive confinement is totally opposite to the canonical front-to-rear polarization of the same factors in directional, adhesion-dependent mesenchymal migration.

### Rear-to-front gradients of EGFR and Rac1 activities are required for leader bleb stabilization

We next examined the requirement for the observed rear-to-front gradient of EGF activity and downstream signaling in maintaining leader bleb stability during migration in non-adhesive confinement. To approach this, we first asked whether the degree of EGFR polarization correlated with bleb morphology or stability. We analyzed bleb aspect ratio as a function of EGFR-GFP polarity index along the bleb for either transient elongated blebs or stabilized leader blebs (Figure 3A-C). This showed that although bleb aspect ratio did not appear to correlate with EGFR polarity for either transient or stable elongated blebs, stable leader blebs generally exhibited greater EGFR polarity than transient elongated blebs (Figure 3B). Furthermore, we found that even if transient blebs had a high aspect ratio, they did not have a high polarity index (Figure 3B). Direct comparison of EGFR polarity index for transient elongated blebs versus stabilized leader blebs confirmed this, with leader blebs showing significantly higher EGFR polarity than transient blebs (Figure 3C). These results show that a strong rear-to-front gradient of EGFR is associated with leader blebs, and suggests that this gradient may be required for their stabilization.

**Figure 3:**
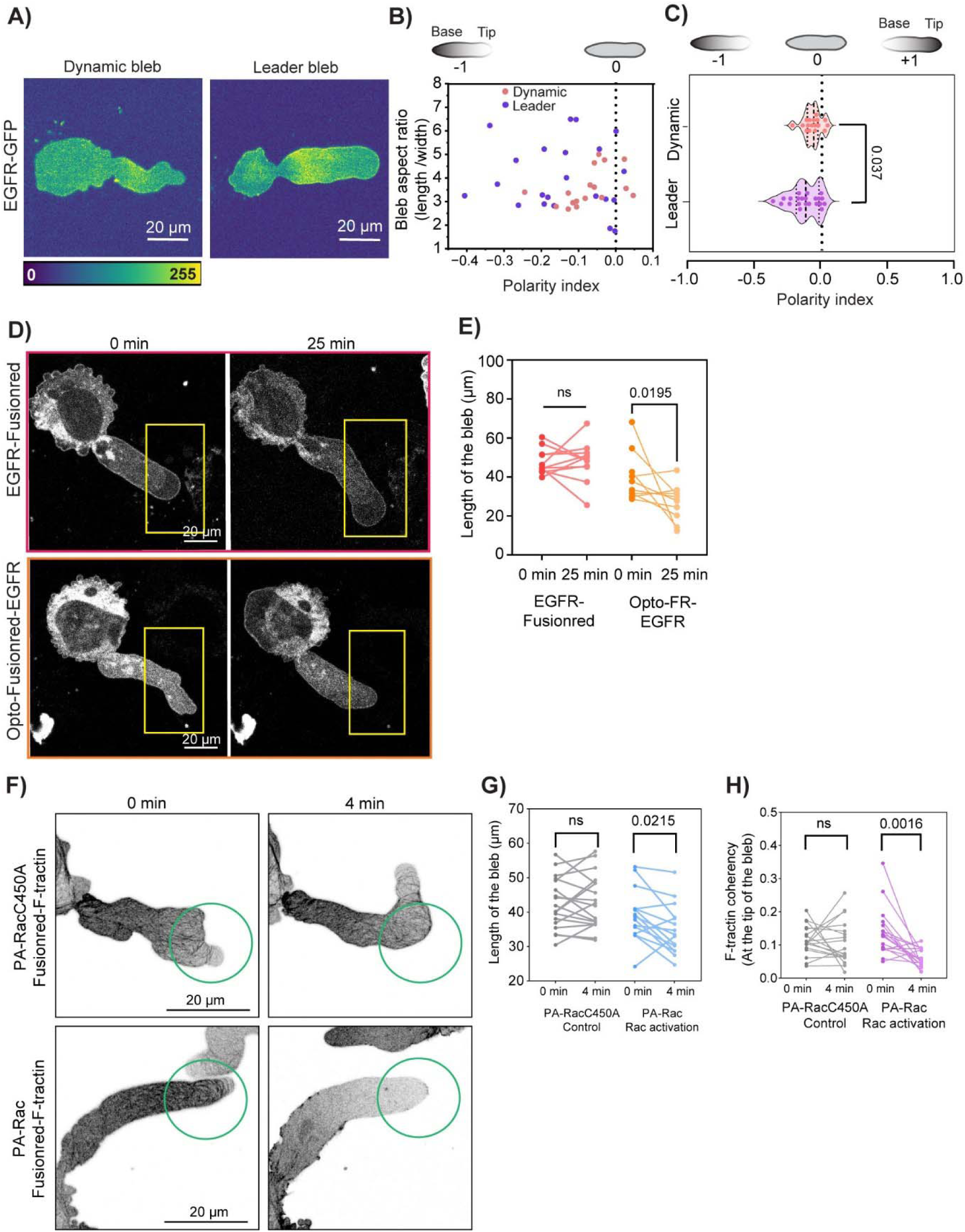
EGFR and Rac1 activity at the rear of the bleb is required for bleb stability. (A) Pseudo colored intensity confocal image of expressed EGFR-GFP in A375M2 melanoma cells under 3μm confinement exhibiting dynamic elongated bleb (left) or leader bleb (right) morphology. (B) Scatter plot of bleb aspect ratio of dynamic elongate blebs (rose, n=17 blebs) and leader blebs (purple, n=22 blebs) blebs versus polarity index of EGFR-GFP in these blebs. (C) Violin plot of polarity indices of EGFR-GFP in both morphologies shown in (A), p=0.037 Welch’s t-test. (D) Stable expression of EGFR-FusionRed (red box, top) and Opto-EGFR (orange box, bottom). Bleb morphology before (left panel) and after continuous illumination (for 25 min, right panel) in the tip of the bleb. Yellow box mark the position of activation illumination region. (E) Quantification and pairwise distribution of the length of the bleb. Each data point represents one leader bleb before and after illumination. Length of the bleb in EGFR-FusionRed construct (n=7 blebs) before (red points) and after 25 min activation (pink points), p=n.s. and Opto-EGFR (n=8 blebs) is before (dark orange) and after 25 min activation (light orange), p=0.0195, Wilcoxon test. (F) Inverted contrast images of FusionRed-ftractin in cells co-expressing either PA-RacC450A (top) or PA-Rac (bottom). Bleb dynamics were imaged before (left) and after continuous illumination for 4 min (right). Green circle marks the region of photoactivation. (G) Quantification and pairwise distribution of bleb length after continuous activation illumination. Length of bleb in cells expressing PA-RacC450A (n=17 blebs) before (dark-gray) and after 4 min activation (light-gray), p=0.0215, Wilcoxon test. PA-Rac (n=14 blebs) before activation (dark-blue) and after activation (light-blue). Paired points represent measurements within one bleb. (H) Quantification and pairwise distribution of F-tractin coherency before and after of same cells in (G). Actin coherency in PA-RacC450A before activation (dark gray) and after activation (light gray). PA-Rac before activation (dark purple) and after activation (light purple); p=0.0016, Wilcoxon test.

To determine the requirement for a rear-to-front EGFR gradient in stabilization of leader blebs, we employed an optogenetic approach to change the localization of EGFR signaling within single cells during migration in non-adhesive confinement. We utilized Opto-EGFR, which consists of fusions between the cytosolic domain of EGFR, a myristoylation-based membrane localization tag, the FusionRed fluorescent protein, the intrinsically disordered domain of FUS, and the photolyase homology domain of *Arabidopsis thaliana* Cry2 to allow reversible, blue light-induced clustering and trans-phosphorylation of the cytosolic domains to mediate spatial control of EGFR signaling ^56,57^. Cells were transfected with either Opto-EGFR or EGFR fused to fusion red (EGFR-FR) and a galvanometric mirror was used to locally expose a 25 × 40 μm^2^ area of the leader bleb tip to repeated pulses every 2 sec of 405nm laser light to locally induce clustering of Opto-EGFR (Figure 3D).Time-lapse imaging showed that light exposure on the bleb tip in cells expressing Opto-EGFR caused a significant decrease in leader bleb length after 25 min compared to similar light exposure on the bleb tip of cells expressing EGFR-FR (Figure 3E). These results show that mis-localizing EGFR signaling to the tip of leader blebs causes bleb retraction, and strongly support the notion that the existing rear-to-front gradient of EGFR activity is required for leader bleb stability.

We then asked whether the asymmetric distribution of Rac1 activity in the rear of the bleb is required for leader bleb stability. We utilized an optogenetic Rac1 construct (PA-Rac) which consists of Rac1 fused to the photoreactive light oxygen voltage (LOV) domain from phototropin ^58,59^, such that 488nm light relieves LOV domain steric hindrance to allow Rac1 activation ^60^. Repeatedly delivering 488 nm light pulses every 4 sec to a 5 μm diameter circle near the tip of the bleb in cells co-expressing PA-Rac and FusionRed F-tractin (F-tractin-FR) was sufficient to decrease bleb length (Figure 3G), cause local disassembly of the actin bundles that reduced actin coherence at the bleb tip (Fig 3F, 3G, Movie 5), drive assembly of lamellipodial actin protrusions (Figure 3F), and eventually retract the blebs (Figure 3G). This was compared to the lack of actin remodeling or retraction in response to similar light exposure at bleb tips in cells expressing the non-photoactivatable control (PA-Rac450A) (Figure 3F-H). Together, these results suggests that spontaneous polarization of EGFR localizes downstream signaling to activate Rac1 at the rear of the leader bleb, and this asymmetric distribution of signaling is required for bleb stability to drive persistent migration in non-adhesive confinement.

### Rear-to-front EGFR gradient is maintained by local restriction of EGFR mobility near the bleb base

We next approached the question of the physical mechanism by which the rear-to-front gradient of EGFR is formed and stabilized within leader blebs in cells migrating in non-adhesive confinement. Maintenance of an asymmetric distribution of trans-membrane proteins that can diffuse in a lipid bilayer could be controlled in several possible ways, including: Localized protein turnover within the membrane by spatially restricted insertion and/or removal by exo- and endocytosis ^61,62^; advective movement driven by direct or indirect interaction with the flow of the underlying cortical actin cytoskeleton ^37,63^ ; or local restriction of free mobility (diffusion and bulk membrane flow) via membrane “corralling” ^64^. We began by ruling out the first hypothesis, which is supported by the fact that leader bleb cells under confinement have their secretory and endocytic apparati localized almost exclusively in the cell body and do not extend into the bleb ^65,66^. We further validated this assumption by photobleaching EGFR-GFP throughout the entire bleb and analyzing the recovery pattern by time-lapse imaging (Extended Figure 2A). This showed that recovery was very slow (Extended Figure 2B) and occurred directionally from the cell body (Extended Figure 2C, Movie 6), with no localized bursts of EGFR-GFP anywhere along the length of the bleb that would be indicative of exocytosis of EGFR-containing vesicles^61,62^.

To differentiate between the hypotheses of cytoskeletal advection and local restriction of free mobility as possible mechanisms of maintaining a rear-to-front gradient of EGFR in leader blebs, we used a combination of mathematical modelling and experiment. The model was based on the assertion that EGFR and membrane synthesis occur in the cell body and thus there is a net movement of protein from the cell body into the bleb.

We considered that the bleb has a geometry based on average measurements (Extended Figure 2D, E), EGFR has an apparent diffusivity ((*v_p_*), Extended Figure 2F, G), a forward advection velocity (*v_p_*), possibly negative retrograde cytoskeletal flow from bleb tip to base (*v_f_*), and a turnover (removal) frequency (membrane *k_m_* and protein *k_p_*), each of which can potentially vary with position along the length of the bleb (L) (Figure 4A).

**Figure 4:**
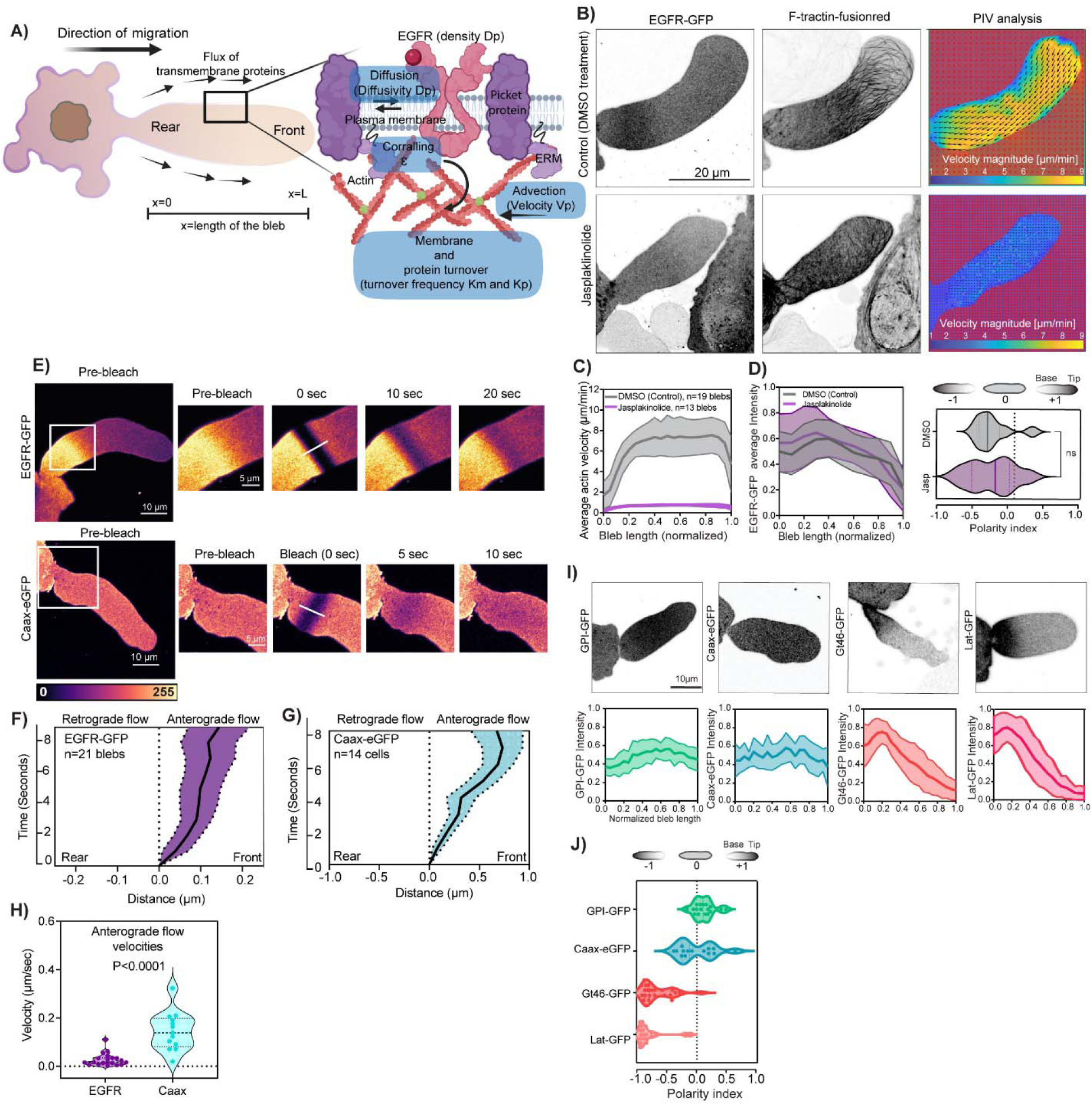
Experiment and modeling indicate neither cortical nor membrane flow mediate EGFR gradient formation in leader blebs. (A) Schematic diagram of a confined cell undergoing leader bleb-based migration (left), and components of the plasma membrane (right). Direction of migration (top right large arrow) and direction of transmembrane protein flux (small arrows around bleb) are highlighted on left panel. Left: EGFR (transmembrane protein receptor, pink) with its ligand EGF (red), picket protein (purple), linker protein (ERM, light purple) that links actin cytoskeleton (red) to the plasma membrane (gray) are highlighted on right panel. Canonical mechanisms for receptor segregation on the plasma membrane including diffusion (Diffusivity Dp), advection (velocity Vp), corralling (corralling ε), membrane and protein turnover (turnover frequency Km and Kp) are in light blue bubbles. (B, E, H) Representative confocal images of blebs in A375M2 melanoma cells under 3μm confinement. B) Left and center columns: Inverted contrast images of blebs in cells co-expressing EGFR-GFP (left) and F-tractin-FusionRed (center), treated with vehicle (DMSO, top) or jaspklinolide (100nM, bottom). Right column: PIV analysis of time-lapse movies of F-tractin-fusionred of the blebs shown at left showing the direction (arrows) and magnitude (color encoded) of actin retrograde flow in two consecutive frames of the movie. (C) Average velocity (dark lines) +/- standard deviation (light lines and shaded areas) of actin retrograde flow in blebs of vehicle (DMSO, gray, n=19 cells) or jasplakinolide (100nM, magenta, n=13 cells) treated cells, normalized by position along the normalized bleb length from N=3 experiments (D) Left panel: Average normalized intensity (dark lines) +/- standard deviation (light lines and shaded areas) of EGFR-GFP in vehicle (DMSO, gray, n=19 cells) or jasplakinolide (100nM, magenta, n=13 cells) treated cells as a function of normalized position along the normalized bleb length. Right panel: Violin plot of polarity indices of EGFR-GFP in blebs of the cells analyzed at left, p=n.s; non-significant by Welch’s t-test. (E) Pseudocolored intensity time-lapse confocal images of a bleb in a cell expressing EGFR-GFP (upper panel) and CAAX-eGFP (lower panel). A small rectangular area was photobleached (at 0 sec) and the fluorescence recovery imaged over time. White line highlights location of intensity of analysis. Mean position of the minima of fitted inverted gaussians relative to the minima position at the time of photobleaching in (F) EGFR-GFP, n=21 blebs; (G) CAAX-eGFP, n=14 blebs. (H) Violin plot of anterograde flow velocity measured from (E,F,G). Each dot represent velocity measured from one bleb. (I) Top row: Inverted contrast image of cells expressing GPI-GFP (left), CAAX-GFP (left center), Gt46-GFP (right center) or Lat-GFP (right). Bottom row: Average normalized intensity (dark lines) +/- standard deviation (light lines and shaded areas) of the probe above each panel as a function of normalized position along the normalized bleb length. (J) Violin plot of polarity indices measured in (I). GPI-GFP (n=17 cells, light green), Caax-GFP (n=14 cells, blue), Gt46-GFP (n=23 cells, dark orange) and Lat-GFP (n=20 cells, Pink).

An important insight gained from initial consideration that EGFR and membrane synthesis occur the cell body is that both bulk membrane and EGFR must both undergo a net anterograde movement from the bleb base towards the tip (See Supplementary text). This was surprising, and in opposition to the notion that advective movement via retrograde cytoskeletal flow would mediate the formation of the rear-to-front EGFR gradient.

We thus explored the possible contribution of advection by actin retrograde flow in forming the EGFR gradient^35^. We first characterized actin dynamics in leader blebs to obtain parameters for our model. Time-lapse super-resolution confocal microscopy and particle image velocimetry (PIV) analysis of F-tractin-FR (Extended Figure 2H and 2I) showed, similar to previous reports ^19,33^, that actin retrograde flow was slow at the bleb tip (∼-5 μm/min), high in the middle (∼-6.5 μm /min), and slower again at the bleb base (Extended Figure 2J, 2K; Movie 7). Modeling the effect of this actin retrograde flow profile on EFGR distribution produced a rear-to-front gradient of EGFR, similar to the measured distribution. Increasing or decreasing the actin retrograde flow speed changed the slope of the gradient (Extended Figure 2L), suggesting that an EGFR gradient could potentially be established by cytoskeletal advection, as has been suggested before^35^. To test this experimentally, we perfused leader bleb cells expressing F-tractin-FR and EGFR-GFP with the actin stabilizing drug, jasplakinolide, to inhibit the actin treadmilling that drives retrograde flow^19^ (Figure 4B, Movie 8). PIV analysis confirmed that jasplakinolide strongly hindered actin retrograde flow along the leader bleb compared to DMSO control (Figure 4B, C). Surprisingly, however, analysis of EGFR localization showed that inhibition of actin flow did not destroy the rear-to-front EGFR gradient, and had no significant effect on the polarity index of EGFR distribution along the bleb (Figure 4D; Movie 8). These results show that although advection by cytoskeletal retrograde flow could in theory mediate formation of a rear-to-front EGFR gradient in leader blebs, experimental evidence shows that this gradient is not established by actin retrograde flow and is independent of its velocity.

We then sought to assess the model prediction related to anterograde membrane and EGFR movement from the cell body towards the bleb tip. To experimentally test this, we expressed either EGFR-GFP (Figure 4E, upper panel) or the membrane marker, GFP fused to the CAAX prenylation signal sequence (CAAX-GFP, Figure 4E, lower panel), photobleached them in a stripe near the base of the bleb, and analyzed the directionality of the intensity recovery by determining the position of the minima of an inverted Gaussian fitted to the bleach stripe over time (Extended Figure 2M and N). This analysis showed that photobleached stripes in either EGFR-GFP or CAAX-GFP exhibited anterograde movement from the bleb base towards the tip (Figure 4F and 4G; Movie 9), but with different apparent velocities with CAAX-GFP moving significantly faster than EGFR-GFP (Figure 4H). Taken together, these results support the notion that EGFR present in the bleb plasma membrane originates by a net anterograde movement from the cell body towards the bleb tip, and that EGFR movement is somehow slowed relative to that of the bulk membrane.

We then explored what type of membrane association caused EGFR to resist anterograde membrane flow and accumulate at the bleb rear to form a rear-to-front gradient. We analyzed the distribution of different types of membrane-associated markers within the leader bleb, including other transmembrane proteins (Gt46-GFP or LAT-GFP (an artificial secretory protein containing a signal sequence, GFP, a consensus N-glycosylation site, the transmembrane domain of the LDL receptor, and the cytoplasmic tail of CD46), an inner leaflet-associated marker (CAAX-GFP), or an outer leaflet-associated marker (GFP fused to glycosylphosphatidylinositol, GPI-GFP) ^67^. Imaging and analysis of polarity index of these markers in leader blebs showed that, similar to EGFR-GFP, the transmembrane proteins LAT-GFP and Gt46-GFP formed rear-to-front gradients, accumulating towards the base of the bleb. In contrast, the inner and outer leaflet membrane markers CAAX-GFP and GPI-GFP were evenly distributed along the bleb (Figure 4I, 4J). These results suggest that transmembrane proteins are capable of resisting anterograde membrane flow to mediate their accumulation in rear-to-front gradients along the leader bleb.

We next pursued the question of how transmembrane proteins resist anterograde membrane flow. We modeled the effect of restriction of EGFR mobility relative to that of the bulk membrane by “corralling” (*E_1_(x)*) but, while considering minimal advection by retrograde actin flow (*hence E_2_(x) = 0.1*), in line with our experimental observations. Our model showed that implementing different corralling and hence different EGFR restriction levels in the leader bleb membrane resulted in rear-to front EGFR gradients that decreased steepness with decreasing corralling (Figure 5A). This suggests that local restriction of EGFR mobility could allow the protein to resist anterograde membrane flow and cause its accumulation in a rear-to-front gradient.

**Figure 5:**
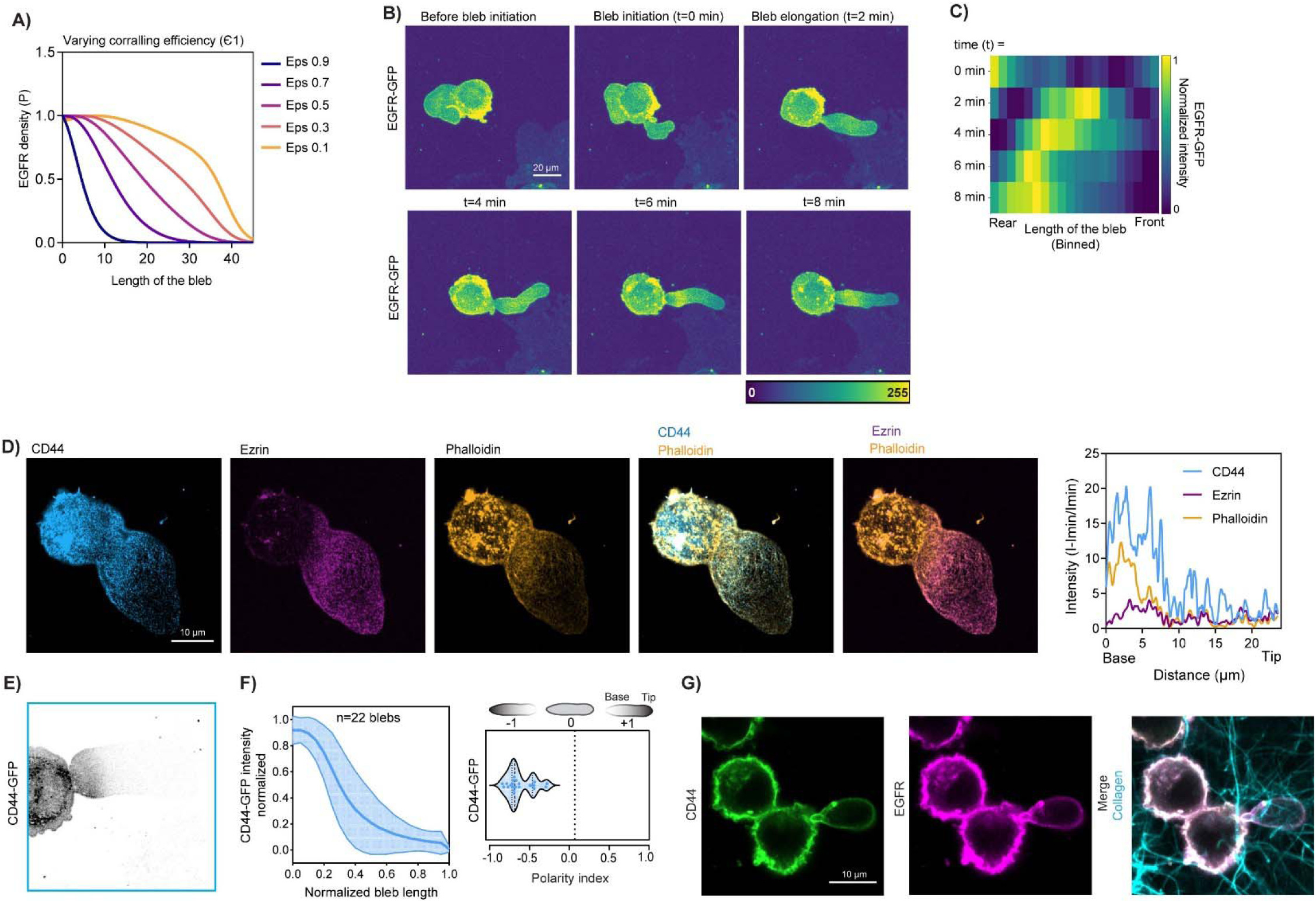
EGFR has restricted diffusion at the rear of the leader bleb. (A) Outcomes of transmembrane protein gradient model. EGFR receptor density as a function of the length of the bleb under different corralling efficiencies ε. Representative confocal images of blebs in A375M2 melanoma cells under 3μm confinement (B-D) or migrating in a 3D fibrillar collagen ECM (G). (B) Left: Pseudocolored intensity time-lapse confocal images of leader bleb formation in a cell expressing EGFR-GFP as the bleb initiates and elongates over time (C) Heat map of the normalized EGFR-GFP intensity of the bleb the rear-to-front over time as a function of normalized length of bleb (binned). (D) Left panels: Immunostaining of CD44 (blue), ezrin (purple) and phalloidin staining of f-actin (orange) and merged images of CD44 phalloidin with Ezrin and Ezrin with phalloidin. Right Panel: Intensity line scan analysis across the leader bleb at left. (E) Inverted contrast image of a cell transfected with CD44-GFP. (F) Graph showing the mean normalized intensity of CD44-GFP (blue box) across the normalized length of the leader bleb, mean ± s.d. (left) and violin plot of CD44-GFP polarity index (right); n=22 cells. (G) Immunostaining of CD44 (green), EGFR (purple) and collagen, (cyan), and merged image of both channels of an A375M cell migrating in a 3D collagen ECM.

To test this prediction, we analyzed the evolution of EGFR-GFP distribution during initial formation of an elongated bleb. We found that as the bleb elongated, EGFR-GFP accumulated at the base of the bleb and never advanced towards the bleb tip, suggesting that EGFR-GFP was corralled at the rear (Figure 5B, 5C; Movie 10). Together, these results suggest that a rear-to-front gradient of EGFR along leader blebs is generated and maintained by restriction of transmembrane protein mobility to mediate bleb stabilization and drive persistent migration in non-adhesive confinement.

### CD44, Ezrin, and a membrane-apposed actin meshwork restrict EGFR mobility at the bleb base to establish the rear-to-front EGFR gradient in leader blebs

We next sought to determine the molecular basis of mobility restriction of EGFR to maintain the EGFR gradient along leader blebs. As CD44 and ERM (Ezrin-Radixin-Moesin) proteins are known to link to the sub-membrane cortical actin meshwork to create membrane corrals that restrict free diffusion of transmembrane proteins in other cell types ^43^, we examined their distributions in leader blebs. Immunostaining for CD44 and ezrin in confined cells revealed a concentration of CD44 and an absence of ezrin in the cell body, but both proteins formed rear-to-front gradients within leader blebs (Figure 5D). Similarly, leader blebs in cells expressing CD44-GFP (Figure 5E) in non-adhesive confinement showed an asymmetric localization with a high concentration at the rear of the bleb that decreased towards the tip (Figure 5F). To determine if cells undergoing bleb-based migration in more physiologically-relevant confined microenvironments exhibited similar protein organization, we assessed by immunostaining the distribution of CD44 and EGFR in melanoma cells migrating in a 3D fluorescently-labeled collagen fibril matrix. 3D confocal imaging showed that in collagen matrices, melanoma cells formed leader blebs in which CD44 and EGFR formed rear-to-front gradients (Figure 5G). Thus, the corralling proteins CD44 and Ezrin localize in a similar rear-to-front gradient as EGFR in leader blebs in cells undergoing confined migration.

We next examined whether actin organization was conducive to corral formation in leader blebs by analyzing actin architecture and proximity to the bleb membrane. Analysis of actin filament organization showed that actin coherence was low towards the bleb base, indicating a dense actin meshwork, and increased towards the bleb tip where filaments formed bundles (Extended Figure 3A, 3B). To determine if the actin meshwork was in close proximity to the plasma membrane, we utilized a biosensor consisting of the actin-binding domain of inositol 1,4,5-Trisphosphate 3-Kinase A fused to citrine fluorescent protein and the CAAX prenylation signal sequence (MPAct-citrine) such that the sensor diffuses freely in the membrane except in regions where cortical actin is within ∼10 nm from the plasma membrane, where it becomes immobilized and enriched by binding actin, thus marking membrane-proximal actin ^68^. Analysis of MPAct-citrine within leader blebs revealed that membrane-proximal actin was predominantly located at the rear of the bleb (Extended Figure 3C, 3D).

Together, these results show that CD44 and Ezrin are concentrated at the rear of leader blebs where the actin cortex forms a dense meshwork in close proximity to the membrane, suggesting these components could form membrane corrals that locally restrict the mobility of EGFR in cells migrating in confined microenvironments.

We next tested the requirement for CD44 and ERM proteins in restricting EGFR mobility by cytoskeletal corralling to maintain the EGFR rear-to-front gradient in leader blebs. We first tested if EGFR mobility correlated with CD44 localization. We performed FRAP to simultaneously bleach small regions near both the bleb base where CD44 concentration is high and at the bleb tip where CD44 is largely excluded, and analyzed their intensity recoveries over time (Figure 6A; Movie 11). This showed that EGFR-GFP intensity recovery at the bleb base was significantly slower than at the bleb tip, indicating a locally reduced mobility of EGFR in the region of CD44 concentration (Figure 6B, 6C). We then tested the requirement for CD44 in forming the EGFR gradient and restricting its mobility in the plasma membrane. Cells were co-transfected with EGFR-GFP and either CD44 pooled siRNAs (CD44-KD) or non-targeting (NT) control and subjected to non-adhesive confinement (Figure 6D). Analysis showed that CD44-KD caused flattening of the EGFR-GFP rear-to-front gradient and significantly skewed the polarity index towards zero compared to NT control (Figure 6E). FRAP experiments and analysis for spots bleached near the bleb base showed that reducing CD44 levels caused significantly faster recovery compared to NT control, confirming that CD44 reduces the mobility of EGFR-GFP (Figure 6F, 6G and 6H; Movie 12). We next tested the involvement of CD44’s link to the extracellular glycocalyx in maintaining the EGFR gradient by treating cells expressing EGFR-GFP in non-adhesive confinement with hyaluronidase (HAase) to degrade polyhyaluronic acid. We found that, compared to DMSO controls, hyaluronidase treatment had no significant impact on the EGFR-GFP rear-to-front gradient or polarity index in leader blebs (Extended Figure 4A and 4B). As A375M2 cells predominantly express ezrin and moesin ^15,69^, we silenced their expression with pooled siRNAs (EM-KD) and assessed the effects on EGFR distribution. This showed that, like CD44-KD, EM-KD flattened the EGFR-GFP gradient along the leader bleb and significantly skewed the polarity index towards zero (Extended Figure 4C, 4D). Importantly, neither CD44-KD nor ezrin-KD caused changes to the organization of actin at the rear of the bleb (Extended Figure 4E and 4F). Thus, CD44 and ERM proteins are required for restricting the mobility of EGFR near the bleb base to maintain the EGFR back-to-front gradient during leader bleb-based migration.

**Figure 6:**
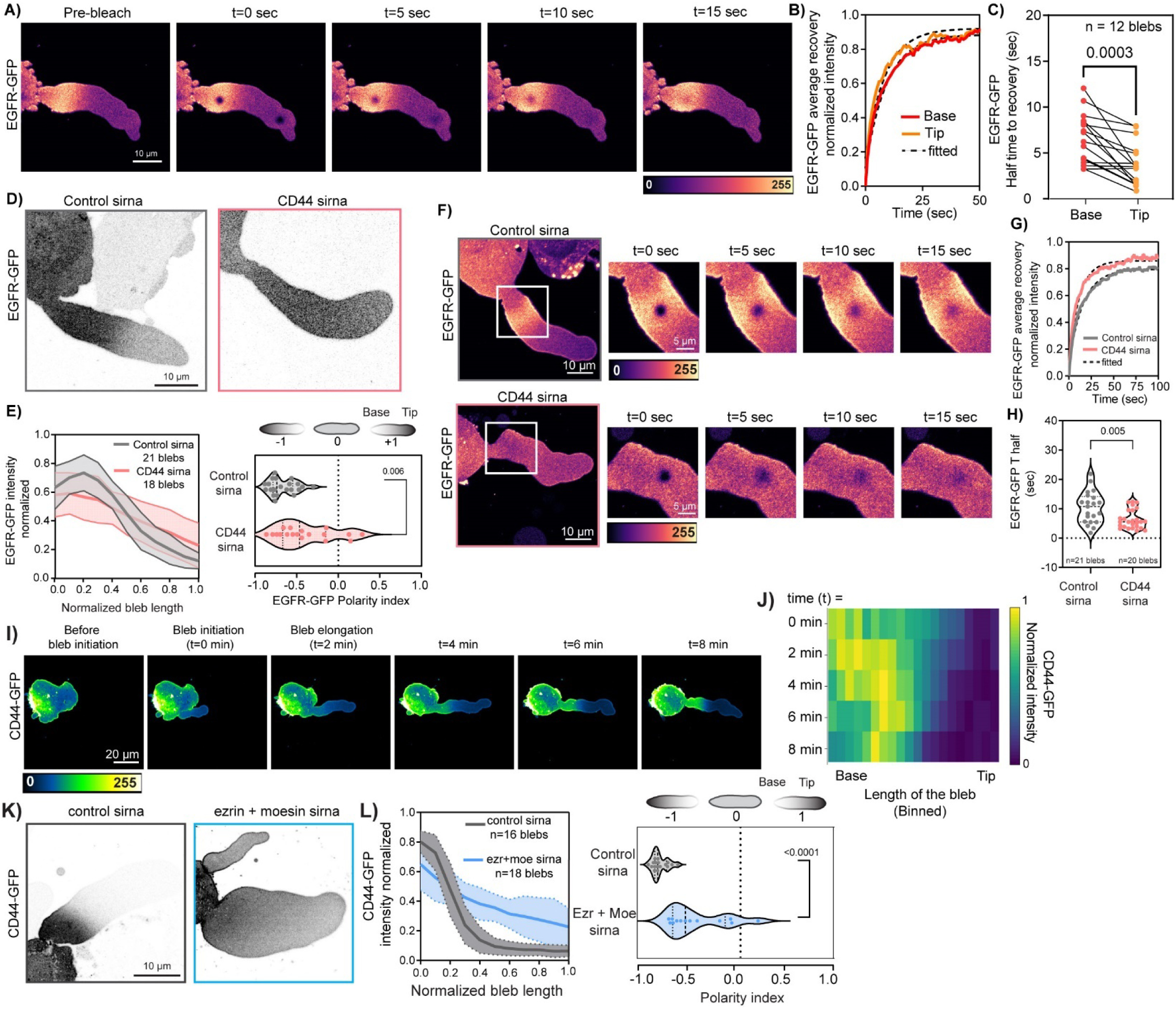
CD44 restricts the diffusion of EGFR at the rear of the leader bleb. (A) Pseudocolored intensity time-lapse confocal images of a leader bleb in a cell expressing EGFR-GFP, with photo-bleaching in two small regions near the front and rear of the bleb. (B) Average normalized fluorescence intensity over time (solid lines) from the two photobleached regions in the cell in the left panel, time zero is the first image after photobleaching. Single exponential least squares fits (dotted lines). (C) Half time of fluorescence recovery after photobleaching determined from single exponential fits of data like that shown in the middle panel, paired points are data from the bleb base and tip within the same cell, n=16 blebs; p=0.0003, paired t-test. (D,F,K) Representative confocal images of blebs in A375M2 melanoma cells under 3μm confinement. (D) Inverted contrast image of Blebs in cells expressing EGFR-GFP and either scramble control siRNA (left) or siRNA targeting CD44 (E) Left panel: Mean normalized intensity profile of EGFR-GFP across the normalized length of the leader bleb, mean ± s.d., from linescan analysis of cells under treatments shown in (D). Right panel: Violin plot of EGFR-GFP polarity index (control-siRNA, gray, n=21 cells, CD44-siRNA, pink, n=18 cells). p=0.006 Mann-Whitney test. (F) Pseudocolored intensity time-lapse confocal images of a leader bleb in a cell expressing EGFR-GFP together with either scrambled siRNA (upper row, Control) or siRNAs targeting CD44 (lower row), with photo-bleaching in a small region near the rear of the bleb. (G) Average normalized fluorescence intensity over time (solid lines) from the photobleached regions in the cells in the left panels, time zero is the first image after photobleaching. Single exponential least squares fits (dotted lines). (H) Violin plot of the half times of fluorescence recovery after photobleaching determined from single exponential fits of data like that shown in (G), control-siRNA, n= 21 cells and CD44-siRNA, n=20 cells, p=0.005 Mann-Whitney test. (I) Pseudocolored intensity time-lapse images of leader bleb formation in a cell expressing CD44-GFP as the bleb initiates and elongates over time (left). (J) Heat map of the normalized CD44-GFP intensity of the bleb the rear-to-front over time as a function of normalized length of bleb (binned). (K) Inverted contrast image of blebs in cells expressing CD44-GFP and either scramble control siRNA (left) or siRNA targeting ezrin (L) Mean normalized intensity profile of EGFR-GFP across the normalized length of the leader bleb, mean ± s.d., from linescan analysis of cells under treatments shown in (K),control-sirna (gray, n = 16 cells) or ezrin and moesin-siRNA (blue, n= 18 cells); N=3 independent experiments. Right panel: Violin plot of CD44-GFP polarity indices for the conditions in (K,L), p<0.0001 Mann-Whitney test.

We next asked how CD44 is maintained in a polarized distribution to restrict EGFR mobility at the bleb base. - Examining the evolution of CD44 localization during leader bleb formation (Figure 6I) showed that, like EGFR, CD44-GFP accumulated at the base of the bleb and never advanced towards the bleb tip as the bleb elongated, suggesting it was trapped towards the bleb rear (Figure 6J; Movie 13). We then probed the role of ERM proteins in CD44 localization by examining CD44 distribution in leader blebs of EM-KD cells (Figure 6K). This showed that EM-KD altered the distribution of CD44 in leader blebs compared to non-targeting controls (Figure 6K and 6L), causing flattening of the CD44-GFP intensity gradient and significantly skewing the CD44 polarity index towards zero (Figure 6L). Taken together, our data demonstrates that ERM proteins promote the polarized distribution of CD44, and together with a membrane-proximal actin meshwork at the rear of the bleb, curtail EGFR mobility as the bleb elongates and to establish a rear-to-front EGFR gradient in leader blebs.

### CD44 is required for polarization of signaling to promote leader bleb stability and persistent migration in non-adhesive confinement

We finally sought to investigate the importance of CD44-mediated corralling of EGFR mobility at the bleb base in promoting bleb polarity and persistent migration in non-adhesive confinement. We first examined the impact of CD44 depletion on polarity signaling within leader blebs. Analysis of Pak-GBD-YFP in CD44-KD cells showed reduction in the rear-to-front polarization of Rac1 activity in CD44-KD compared to NT controls (Figure 7A and 7B). To assess the role of CD44 on leader bleb-based migration we silenced CD44 expression by siRNAs targeting the 3’UTR of CD44 (CD44-3’KD) and compared this to NT controls. This showed that CD44-3’KD significantly inhibited leader bleb formation (Figure 7C and 7D; Movie 14) and drastically reduced the migratory range of cells in non-adhesive confinement (Extended Figure 5A). However, cells that managed to undergo leader bleb-based migration exhibited persistent migration with similar velocity (Figure 7E and Extended Figure 5B; Movie 15). Notably, reintroducing CD44-GFP into the knockdown cells rescued the percentage of cells undergoing leader bleb-based migration (Figure 7D) and restored the migratory range and persistence to levels comparable to NT controls (Figure 7E, Extended Figure 5A). Together, our results show that during non-adhesive confinement, melanoma cells protrude elongated blebs that trap CD44 at their base due to its interaction with the underlying actin cortex via ezrin, that in turn restricts EGFR mobility to mediate polarized signaling within the bleb to stabilize bleb morphology and promote persistent leader-bleb based migration (Extended Figure 5C).

**Figure 7:**
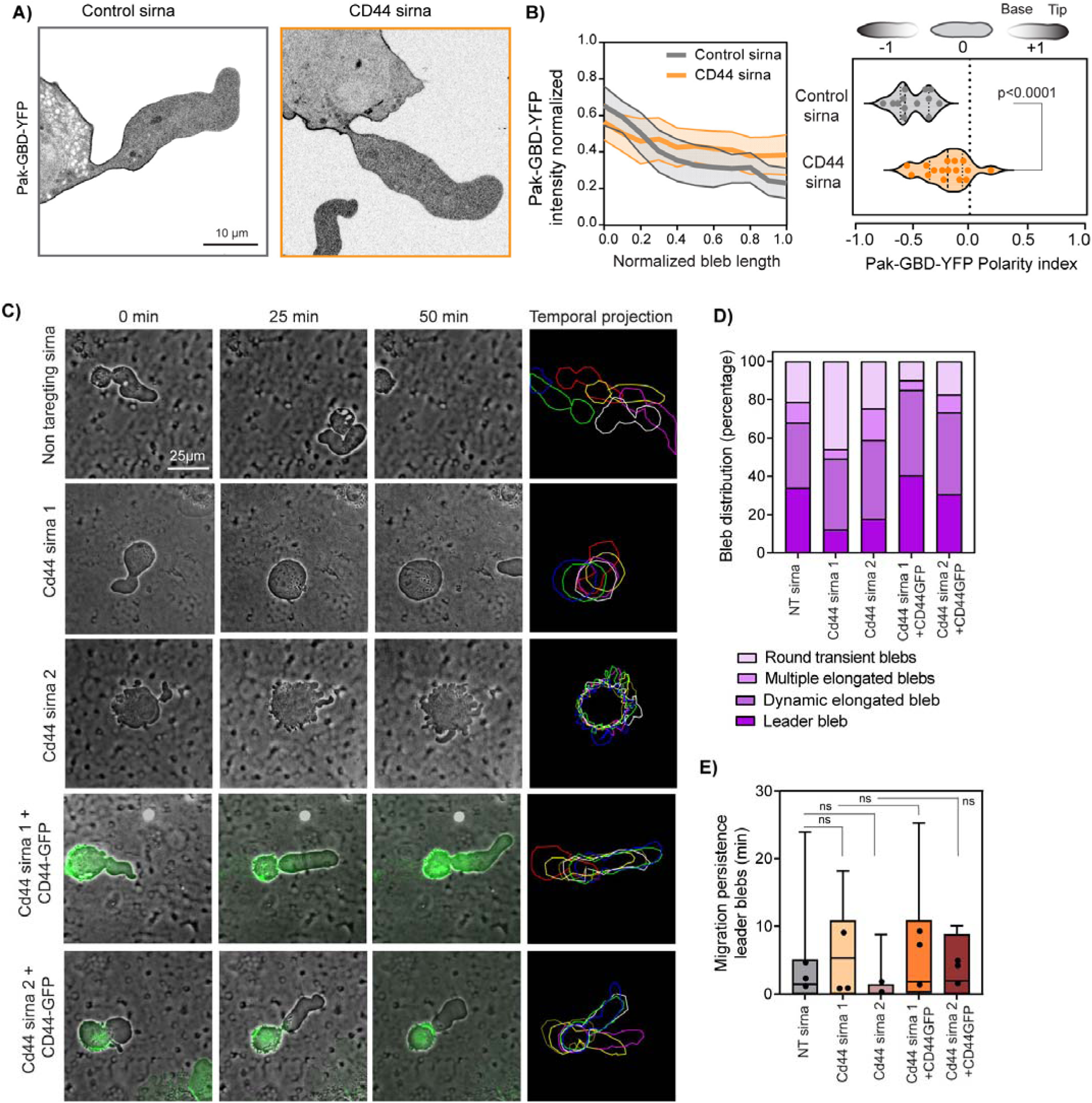
CD44 is required for rear-to-front Rac1 activity gradient and leader bleb stability and persistent migration under confinement. (A) Inverted contrast confocal images of cells co-expressing Pak-GBD-YFP to localize active Rac1 and either control-siRNA or CD44-siRNA, (B) Left panel: Mean normalized intensity profile of Pak-GBD-YFP across the normalized length of the leader bleb, mean ± s.d., from linescan analysis of cells under treatments shown in (A). Right panel: Violin plot of Pak-GBD-YFP polarity. Control-siRNA (n=13 cells), CD44-siRNA (n=16 cells) (C) Representative time-lapse phase-contrast images (left three columns) of A375M2 melanoma cells under 3 µm confinement and temporal color-encoded projection of cell shapes at 10-minute intervals (right column) of the cells shown at left. Cells were transfected with either non-targeting Control-siRNA (top row) or siRNAs targeting two different sequences in the 3’-UTR of CD44-siRNA either alone (sequence 1 (second row) or sequence 2 (third row)) or together with CD44-GFP (green, sequence 1 (fourth row) or sequence 2 (bottom row)) (D) Quantification of bleb phenotype distribution percentage for the conditions in (C). (Control-sirna n=47 cells, CD44-siRNA1 n=122 cells, CD44siRNA2 n=69 cells, CD44-siRNA1 + CD44-GFP n=121 cells, CD44siRNA2 + CD44-GFP n=75 cells) (E) Box plots of migration persistence of cells with leader blebs for the conditions in (E) (Control-sirna n=25 cells, CD44-siRNA1 n=10 cells, CD44siRNA2 n=12 cells, CD44-siRNA1 + CD44-GFP n=20 cells, CD44siRNA2 + CD44-GFP n=17 cells). Box plot showing the median value and the box extending from the 25^th^ to 75^th^ percentiles and whiskers show the minimum and the maximum values of the data. P value, not significant (ns), Welch’s t-test, Scale bar= 25 microns.

## Discussion

Cells with high contractility under confinement form hydrostatic pressure-driven blebs, enabling efficient migration in low-adhesive environments ^17,70^. Our study reveals the molecular basis for maintaining polarity in metastatic melanoma cells migrating in non-adhesive confinement, showing that confinement-induced pressure promotes a persistent, highly polarized stable leader bleb. This stability is achieved through an asymmetry in actin polymerization at the bleb leading edge and contractility at the bleb rear, coupled with actin retrograde flow, which creates a feedback loop that sustains the polarized morphology in the absence of any external chemical cues or integrin signaling ^19,20,22,71^. We investigated the basis of the polarized bleb morphology of metastatic melanoma cells migrating in confined, non-adhesive environments. Our findings reveal that EGF signaling via PI3K stabilizes and maintains a polarized leader bleb for efficient migration.

Using high-resolution microscopy and biosensors, we observed a unique rear-to-front gradient of EGFR/PI3K activity, promoting PIP3 and Rac1 accumulation at the bleb rear, with its antagonists PIP2 and RhoA concentrated at the bleb tip. Optogenetic experiments showed that mislocalizing EGFR and Rac1 activities causes bleb retraction, underscoring the role of this signaling gradient in bleb stability. Mathematical modeling and experiments identified a mechanism where, as the bleb initiates, CD44 and ERM proteins restrict EGFR mobility in a membrane-apposed cortical actin meshwork in the bleb rear, establishing a rear-to-front EGF-PI3K-Rac gradient. Thus, our study reveals the biophysical and molecular underpinnings of cell polarity in bleb-based migration of metastatic cells in confined microenvironments (Extended Figure 5C).

This study reveals a striking polarization of signaling within leader blebs in metastatic melanoma cells under confinement that is the reverse of the classical model of cell polarity seen in mesenchymal migration. In integrin-dependent mesenchymal migration, ECM engagement activates integrin-PI3K-Rac1 signaling at the leading edge ^72^, promoting Arp2/3 mediated, branched actin polymerization-driven protrusion of lamellipodia ^29^, while Rho GTPase coordinates activation of myosin II contractility in the cell center and rear that drives actin retrograde flow at focal adhesions ^73,74^, acting as a regulatable molecular clutch by mediating transient interactions between actin and ECM-bound integrins to drive cells forward ^13^. By contrast, we observed a reversed polarity in bleb-based migration in non-adhesive confinement, with EGFR-PI3K-Rac signaling localized at the bleb rear and activated RhoA concentrated at the bleb leading edge. How does this *reversed* signaling polarity drive forward movement in leader bleb-based migration? We suggest that the Rho-GTP activity we documented at the bleb tip likely mediates formin-dependent assembly of actin bundles observed at the leading edge, in line with recent work showing that mild formin inhibition halts actin nucleation at the bleb tip and retrograde flow ^33^. However, since leading edge protrusion in bleb-based migration is driven by pressure, the branched actin network that is believed to be evolved for generating membrane pushing forces ^75^ is not needed. But leading edge actin assembly is still required for bleb based migration in order to feed continuous retrograde flow along the bleb length to sweep transmembrane, cytoskeleton-associated proteins rearward to mediate the non-specific friction with the substrate that drives the cell forward ^19^. Why leading edge Rho activity does not promote contractility at the bleb tip is not clear, as myosin II filaments have been shown to assemble at the bleb midpoint and grow into ribbons as they contract and flow rearward with the dense actin meshwork at the bleb rear ^33^. The transition from actin bundles at the bleb tip to meshwork at the bleb rear is likely mediated by the spatial switch between leading edge Rho activity to EGFR-PI3K-Rac1 signaling to activate Arp2/3-mediated actin meshwork assembly at the bleb rear. Furthermore, Rac1 has been shown to drive myosin II filament assembly at focal adhesions in mesenchymal migration ^76^, and in the absence of integrin signaling, may mediate myosin II assembly in the bleb rear to pull actin rearward, disassemble it at the bleb neck, and actively deplete actin from the cell front to maintain blebbing ^19,33^. This spatial reorganization of Rho-GTPases and PI3K signaling may allow metastatic cancer cells to use the same basic motility building blocks in a different organization to shift from adhesion-dependent mesenchymal migration to bleb-based, adhesion-independent migration under confinement, creating distinct types of protrusions suited to varying environments.

Our study identifies CD44 as critical for maintaining polarity and bleb stability in metastatic melanoma cells, in line with its high expression at tumor fronts^38^. CD44 depletion reduces blebbing and contractility, implying a role in myosin activation at the bleb rear^70^. One mechanism may involve ligand-dependent activation, where CD44 binds hyaluronan or matrix metalloproteinases to recruit Rho-GTPases, activating myosin II ^77,78^.

Alternatively, our findings suggest that CD44 acts as a transmembrane “picket”, to corral EGFR, via an ezrin and actin meshwork, at the bleb rear, where lower receptor density at the front limits activation ^79^. This configuration could form a feedback loop as growth factor signaling promotes downstream ERM protein phosphorylation, linking CD44 to actin. Although hyaluronan degradation didn’t alter EGFR polarity in our experiments, this mechanism might vary in the tumor microenvironment. Our data reveal that CD44 restricts EGFR diffusion across the bleb, impacting its spatial distribution. Additionally, changes in the cytoskeletal network during the bleb cycle affect diffusion, with denser actin during bleb retraction aiding myosin II-driven retraction ^33^, potentially engaging more transmembrane “pickets” to redistribute EGFR signaling and regulate bleb dynamics during bleb-based migration.

The spontaneous polarization of metastatic melanoma cells in the absence of chemical or haptotactic gradients, coupled with their efficient migration in confined, low-adhesion environments, has important implications for tumor invasion and metastasis. Previous studies have shown high Erk signaling in melanoma cell blebs, likely due to localized, restricted signaling ^20^. High contractility can increase the invasive potential of cancer cells, confer anoikis resistance ^80,81^, and promote drug resistance ^70^. How common these phenomena are across different solid tumors and how they change with disease progression remain important areas for further exploration.

## Methods

### Cell culture

A375MA2 cells were obtained from American Type Culture Collection (ATCC CRL-3223, ATCC, Manassas, VA) and were maintained at 37°C at 5% CO_2_ for <15 passages in tissue culture treated dishes (Falcon). Cells were maintained in DMEM supplemented with 10% FBS (Atlanta Biologicals), GlutaMAX (Gibco) and 20 mM Hepes pH 7.4. Transient transfection of cDNAs or siRNAs was performed using the Amaxa nucleofector kit V (Lonza) with 1.5 µg of DNA for 2×10^6^ cells, or with 1 µM of siRNA for 48 hours prior to experimental use. For drug treatments or siRNA or cDNA transfections, cells were plated on 6 well pre-treated tissue culture plates (Nunc 6 well multiwell plate, Thermo fischer scientific). For serum starvation, cells were plated on 6 well tissue culture plates for 2 hours and then culture media was replaced with DMEM without FBS for 12 hours. Cells were trypsinized, pelleted and fresh FBS-deprived media was added to the cells. For experiments with EGF stimulation in serum-starved cells, media supplemented with 50ng/mL EGF (Thermo scientific) was added last.

To culture melanoma cells in 3D collagen ECM, 2.5 mg/ml Type 1 collagen was mixed with A375MA2 cells. Briefly, 35 mm glass bottom dishes were plasma cleaned and then 300µl of 1.5 mg/ml collagen mixed with 3000 cells were gently pipetted on the glass bottom dish to make a thin layer and was allowed to polymerize at 37°C for 1 hour. Following polymerization, 500µl of 1.5 mg/ml collagen was added on the top of the collagen with cells to secure it to the bottom of the dish for imaging. Collagen was again allowed to polymerize for another hour at 37°C. Upon final polymerization, 1 ml of culture media was added gently to the dish and left at 37°C overnight. Fresh culture media was added to the dish an hour before imaging or fixation.

### cDNA expression vectors and lentiviral expression

The following cDNAs were used for DNA transfections: EGFR-GFP (Addgene 32751), Akt-PH-GFP (kind gift from Tamas Balla), GFP-C1-PLCdelta_PH (Addgene 21179), Pak-GBD-YFP, Rhotekin-RBD-GFP, (kind gifts from Klaus Hahn), pTriEX-PA-Rac1-C450A (Addgene 22025), pTriEx-PA-Rac1 (Addgene 22024), F-tractin-FusionRed (kind gift from Michelle Baird from Mike Davidson collection), CD44-GFP (kind gift of Satyajit Mayor), GPI-GFP, Gt46-GFP, Lat-GFP, Caax-GFP (kind gifts from Jennifer Lippincott-Schwartz), C1-MPAct-mCitrine (Addgene 155220). EGFR-FusionRed, Myr-FusionRed-Cry2Drop-EGFR and Clover-ZtSH2 and EGFR-CD3ε-FusionRed. ON-TARGET plus SMART pool siRNAs (Dharmacon) were used for human CD44 (L-009999-00-0005), EGFR (L-0003114-00-0005), Ezrin (L-017370-00-0005), siGENOME siRNA Human EGFR (D-003114-34-0005). For CD44 rescue, two independent siGENOME 3’UTR human CD44 siRNA were utilized (Dharmacon, D-009999-20-0005, D-009999-21-0005).

### Lentivirus

Lentiviral particles were generated by transfecting HEK293FT (∼1 million cells) cultured in DMEM with Lipofectamine 2000 (Thermo Fisher) according to manufacturer’s protocols using 2.5 µg plasmid DNA of interest, 1 µg pMD2.G, and 2.5 µg psPax2, and collecting the virus-containing supernatant at 48 hours in media without FBS. Supernatant was filtered through a 0.45 µm filter (Sigma), then immediately placed on target A375M2 cells that had been seeded 24 hours before transduction at a 1:1 ratio of viral supernatant to supplemented DMEM media and allowed to incubate for 48 hours, after which cells were passaged into media containing a final concentration of 2.0 µg/ml puromycin for selection. Stable cell lines were generated with the following constructs: EGFR-FusionRed, Myr-FusionRed-Cry2Drop-EGFR and Clover-ZtSH2 and EGFR-CD3ε-FusionRed ^52,57^, selected with 2.0µg/ml puromycin, and then expanded for 2 weeks or more (to reach around 80 percent confluency), followed by sorting by flow cytometry for Clover- and FusionRed-positive cells of medium fluorescence intensity level.

### Drug Treatments

Working concentration of Erlotinib 10µM (Sigma-Aldrich, SML2156), Wortmannin 10nM (Sigma-Aldrich, W3144) and Jasplakinolide 100nM (Invitrogen), Hyaluronidase 20 units/mL from bovine testes (Sigma-Aldrich)

### Cell confinement assays Dynamic confiner

To subject the cells to confinement for high resolution live cell-imaging, the 1-well Dynamic Cell Confiner System (4Dcell) was used per manufacturer’s directions ^82^. The device consists of a pressure cup, a confiner coverslip affixed with micro-pillars and a glass bottom dish ^19^. The height of the micropillars determined the height for the spatial confinement of the cells between the coverslip and the substrate, which was 3 microns in this study. The confiner coverslips were sonicated in PBS, air dried under the hood. The glass bottom dishes (Fluorodish, 35mm, FD3510-100) and the confiner coverslips were plasma-treated for 1 min, incubated with 0.5 mg/mL pLL-*g*-PEG (SuSoS, PLL(20)-g[3.5]-PEG(2)) in 10 mM pH 7.4 HEPES buffer for 1h at room temperature. Confiner coverslips and glass-bottom dishes were rinsed briefly with PBS and incubated in cell culture medium for at least 2 hours before confining the cells. Prior to confinement, cells were trypsinized (Tyrpsin-EDTA 0,25%), washed, and collected in a pellet by brief centrifugation at 3,000 rpm and resuspended in warm DMEM with FBS (without phenol red) and then plated on glass bottom dishes that were pre-treated with PLL-g-PEG and incubated at 37 °C for 30 min and was then placed on the microscope. The confiner cover slip with pillars attached to the confiner suction cup was then lowered over the sample and attached with low pressure seal sufficient for attachment to the glass dish but insufficient for confinement of the cells. Confinement was initiated slowly by increasing the pressure utilizing the custom software (4Dcell) to control the vacuum pump.

### Agarose pads

Cell confinement for immunostaining was performed under agarose slabs. Agarose slabs for cell confinement ^18^ were made by boiling 750mg of ultrapure agarose (Life Technologies) added to 50 ml of DMEM without phenol red and pouring 1 ml directly in a fluorodish (35mm, FD3510-100). After gelation, a hole was punched in the agarose using a small biopsy needle. Prior to confining, 1 ml of 1% BSA made in DMEM was added to the agarose slabs and was left to equilibrate and coat overnight. Before use, media with BSA was thoroughly aspirated and 100μl of media containing cells were pipetted below the agarose pad by gently lifting the pad slightly from the empty hole. The remaining media was thoroughly vacuumed out of the punch hole. To prevent drying, the dish was sealed using parafilm and was incubated for 1 hour in the incubator. After an hour, the immunostaining was performed using the protocol described below.

### Immunofluorescence

Samples under agarose pad confinement were fixed for 20 min at 37°C with 4% paraformaldehyde (PFA; Electron Microscopy Science,15710) in cytoskeleton buffer (CB; 10 mM MES, 3 mM MgCl2, 138 mM KCl, 2 mM EGTA) and the agarose pad was gently removed. Cells were the carefully permeabilized with 0.5% Triton X-100 in CB at RT for 5 mins, and quenched with 10 mM glycine in CB at RT. Cells were washed 2 × 5 mins then 2 × 10 mins with Tris Buffered Saline (TBS; 20 mM Tris, pH 7.6, 137 mM NaCl2) before blocking for 1 hour at RT with blocking solution (2% BSA IgG free and protease free (Sigma-Aldrich, A3059); 0.1 % Tween 20 (Sigma-Aldrich, P1379) in TBS). Cells were incubated for 2 hours at room temperature (RT) or overnight at 4°C with the following primary antibodies anti-CD44 (clone-BJ18; 338802, Biolegend), anti-Ezrin (3C12, Invitrogen), anti-EGFR (H11, MA5-13070, Invitrogen) in TBS. Dishes were then washed 3X with TBS and incubated with the following fluorophore-conjugated secondary antibodies (1:500), Alexa Fluor 488 Donkey anti-rabbit (711-545-5152, Jackson), Alexa Fluor 594 Donkey anti-rabbit (711-585-152, Jackson), Alexa Fluor 488 Donkey anti-mouse (715-545-150, Jackson) and Alexa Fluor 594 Donkey anti-mouse (715-585-150, Jackson), Alexa-Fluor-647 phalloidin (1:200; ThermoFisher, A22287) diluted in blocking solution, for 1 hour at RT. Cells were washed with TBS (2 × 10 mins, each). Coverslips were mounted on glass bottom dishes in mounting media (Dako; Pathology Products, S3023).

### Immunostaining of cells in 3D collagen ECM

Cells embedded in collagen in 35mm glass-bottom dishes were fixed for 60 min at 37°C with 4% paraformaldehyde in cytoskeleton buffer (CB) (10mM Mes, PH 6.1, 138mM KCl, 3mM MgCl, 2mM) then permeabilized with 0.5% of Triton X-100 diluted in CB at room temperature for 3 hours. The samples were then washed 3 × 15 min with TBST (0.1% Tween-20), then 3 × 10 min with TBS before blocking at 4°C overnight.

For primary antibody staining, collagen samples containing cells were incubated at 4°C overnight with anti-CD44 (clone-BJ18; 338802, Biolegend), anti-Ezrin (3C12, 357300, Thermo Scientific)and anti-EGFR (H11, MA5-13070, Invitrogen) in TBS. Samples were then washed 3X with TBS and incubated with the following fluorophore-conjugated secondary antibodies (1:500), Alexa Fluor 594 Donkey anti-rabbit (711-585-152, Jackson), Alexa Fluor 488 Donkey anti-mouse (715-545-150, Jackson) diluted in blocking solution, for 3 hours at RT. After washing with TBS 3X 10 min each, samples were taken for imaging.

### Immunoblotting

Cells were lysed in 2X SDS sample buffer, separated by SDS -PAGE (4-20% Tris-Glycine gels (Thermo Fisher Scientific) and electro-transferred at room temperature (RT) for 1 hour to Immobilon-P PVDF membrane.

Membranes were blocked for an hour at RT with 5% nonfat dry milk in TBS-T buffer (TBS + 0.1% Tween-20) then incubated for 2 h at RT with indicated primary antibodies. Subsequently, membranes were washed 3 × 5 min in TBS-T, incubated with appropriate HRP-conjugated secondary antibodies (1:10,000) for 1h at RT then washed 3 × 5 min in TBS-T. An ECL detection system (iBright 1500, Invitrogen) was used to visualize protein bands. The following antibodies were used anti-phospho-EGFR (1:1000; Tyr1068, D7A5 Cell signaling technology), anti-EGFR (1:1000; D381B1, Cell signaling technology), anti-phospho-Akt (1:2000; Ser473, D9E, Cell signaling technology), anti-GAPDH (14C10, Cell signaling technology). The secondary antibodies (HRP-conjugated goat anti-mouse (115-035-003) or goat anti-rabbit (111-035-003) were from Jackson ImmunoResearch Laboratories.

### Microscopy

#### Spinning Disk Confocal Imaging

Confocal fluorescence imaging was performed on a Nikon Eclipse Ti2 microscope equipped with a Yokogawa CSU-W1 spinning disk scan-head along with Perfect FocusTM. Either a Nikon Plan Apo 60x oil 1.49 NA DIC or Plan Apo 20x/0.75 Ph2 DM Nikon objective lens was used. Illumination was provided by a Nikon LUNV 6-line laser unit, and images were captured with a Hamamatsu Orca-Flash 4.0 v3 camera. Phase contrast illumination was provided by a 100W halogen bulb or LED using an 0.52 NA condenser lens. Microscopes were equipped with the Nikon motorized stage with xy linear encoders and a Mad City (Madison, WI) Nano-Z100 piezo insert with 200um travel. Laser confocal or DIC illumination were selected with electronic shutters and an automated filter turret containing a multi-bandpass dichromatic mirror together with an electronic emission filter-wheel. Microscope functions were controlled by NIS-Elements software (Nikon). Cells were imaged in 35 mm glass bottom dishes (FluoroDish) or the 1-well Dynamic Cell Confiner System (4Dcell), depending on experimental conditions.

#### Laser Scan Confocal Imaging

Fluorescence recovery after photobleaching (FRAP) was performed on an inverted Nikon A1R resonance-scanning confocal microscope on a Nikon Ti body with perfect focus system (PFS) and a motorized stage with xy encoder and Mad City piezo stage insert. Images were acquired with a Nikon Apo 60x/1.4 oil λS DIC N2 objective lens and controlled by NIS-Elements software (with advanced research package). For photobleaching experiments, the 488 nm or 405 nm Nikon LU-n4 laser was used to photobleached using the Galvano scanner and the same scanner was used for acquisition.

#### Super-resolution Confocal Imaging

Imaging of confined live and fixed cells for super-resolution was performed using a Zeiss LSM880 with the Axio Observer7 confocal microscope stand controlled by Zen black software with piezo-Z stage (Wienecke & Sinske GmrH) equipped with Airyscan, using a Plan-Apo 63x 1.4 NA DIC M27 oil objective with a 32-channel GaAsP-PMT area detector. Airyscan image reconstructions were processed in auto strength mode using ZenBlack software (Version 2.3). 488 nm multi-line 25 mW Argon laser, 561 DPss solid state 15mW laser and 633nm HeNe 5mW lasers were used. For optogenetics experiments, the bleaching sub-module with the 488nm laser was used (using 10-20% laser power) and acquired using multidimensional acquisition module of the Zeiss software. Additional analysis was performed in ImageJ (NIH).

### Image Analysis

#### Cell migration analysis

Migration analysis of the cells under confinement was performed on time-lapse phase contrast image series (acquired every 1min for 60 min) in Fiji using “Manual Tracking plug-in” (MTrackJ; ^83^) where each step was manually tracked following the center of the cell body. The tracking coordinates were then exported to the Microsoft Excel plug-in Diper^84^ and mean squared displacement over time and speed were computed according to:

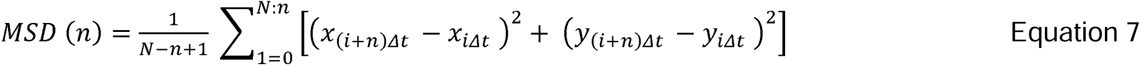

by utilizing overlapping time intervals. MSD is computed for a given cell for step size *n* with N being the total number of displacements per trajectory and *Δt* is the minimum time interval between the adjacent points in the trajectory.

The MSD from each track was utilized and the data was fitted to Furth’s formula according to persistent random walk (PRW) model of migration and is described in equation 8 to obtain persistence time P and diffusion coefficient D.

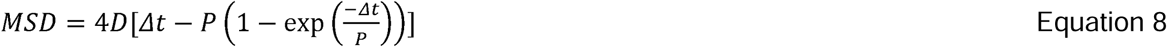

Fittings were performed in Origin (Pro) using the non-linear curve function with orthogonal distance regression. Persistence and diffusion coefficient values were then plotted using GraphPad Prism.

#### Bleb distribution and lifetime

From the phase contrast imaging of cells migrating under confinement, blebbing cells were categorized based on the bleb lifetime. Time duration required for a bleb cycle (nucleation, elongation, persistence and retraction) was noted as the bleb lifetime. Small round blebs with lifetime less than 1 minute were categorized as transient blebs. Other blebs with a lifetime more than 1 minute and which were cylindrical in shape were categorized as elongated blebs. Lengths of the elongated blebs were measured for each time point when the bleb existed in the image frame using the segmented line function in Fiji by drawing regions of interest (ROI) manually. Cells with multiple such blebs were categorized as multiple elongated blebs, and polarized single large blebs were further grouped into dynamic elongated bleb (with a bleb cycle of less than 25 minutes) or leader blebs (with a bleb cycle of more than 25 minutes).

#### Normalized intensity profiles

Intensity profiles of blebs expressing fluorescently tagged membrane proteins were calculated using a segmented line function in Fiji by drawing an ROI of 10 point width from the base of the bleb to the tip of the bleb. Measured intensity was normalized to the maximum intensity and was divided as a function of bleb length into 11 bins, taking 0 for the bleb base and 1 for the bleb tip. Normalized mean intensity profiles from all the blebs were averaged for each position bin and plotted.

#### Polarity index

Polarity index in elongated blebs was defined as PI = (I front-I rear) / (I front + I rear) where I front is the integrated intensity of fluorescently tagged membrane proteins at the bleb tip (at bin 1 of the bleb length) and I rear is intensity at the bleb base (at bin 0 of the bleb length).

#### Analysis of EGFR sensor

Intensity profiles from EGFR FusionRed and EGFR sensor were calculated using a segmented line function in Fiji by drawing an ROI of 10-point width from the base to the tip of the bleb.

Measured intensity was normalized to the maximum intensity and was divided as a function of bleb length into 11 bins, taking 0 for the bleb base and 1 for bleb tip. The ratio of the intensity of EGFR sensor to EGFR was calculated and plotted in a similar fashion. Normalized mean intensity profiles from all the blebs were averaged for each position bin and plotted. Polarity indices of EGFR sensor, FusionRed and ratio were computed and plotted as described above.

#### Bleb aspect ratio

Aspect ratio of elongated blebs was computed by first measuring the length of the bleb and maximum width of the bleb by using a segmented line function in FIJI by drawing a ROI across length and width of the bleb. Second, ratio of length to width were computed and plotted for several blebs.

#### Length of bleb pre and post photo-activation

For EGFR activation in the tips of the blebs, a rectangular ROI was selected for defining the region of activation of size ∼60 × 20 µm for photo-activation in the blebs of the cells expressing Opto-EGFR-FusionRed or EGFR-FusionRed. ROIs were selected such that it covered the tip of the bleb and some additional space for bleb movement. Continuous activation using 488nm laser was performed in this defined region with acquisition approximately every 3 sec for around 25 min. For Rac activation experiments, a circular ROI of diameter ∼15µm was selected for defining the region of activation of blebs in the cells expressing PA-Rac or PA-Rac-450A with F-tractin-FusionRed. Continuous activation using 488nm laser was performed with image acquisition every ∼1 sec for 5 min to capture changes in the actin dynamics and architecture. To measure the bleb length pre- and post-photoactivation, a segmented line function was used in Fiji (ImageJ) to a draw line from the base of the bleb to the tip and values corresponding to the length of the bleb were extracted and exported to graph.

#### Actin coherency measurement

Actin coherency was measured in the cells expressing F-tractin-FusionRed by using the OrientationJ macro in Fiji (ImageJ) that derives local orientation and isotropic values (coherency and energy) of every pixel. Coherency across the whole leader bleb was measured by dividing it into 20 equal oval regions of interest (ROI) slightly overlapping each other from the rear to the tip of the bleb. The area of the ROIs were-based on the area of the leader bleb. The coherency from each ROI was extracted and average was computed and plotted for many blebs.

### Fluorescence Recovery After Photobleaching

#### Spot FRAP and analysis

For spot fluorescence recovery after photobleaching (FRAP) experiments, the intensity recovery over time was extracted from small area of ∼1 micron diameter where photobleaching was performed. Intensity data was extracted from this region pre- and post-bleaching. Two other similar sized ROIs were used to extract intensities from the background of the image and another region in the cell used for bleaching control. The extracted intensities from each ROI was then uploaded in EasyFRAP-web FRAP analysis tool ^85^. Intensity values were normalized and fit with a single term exponential equation. The web tool computes t_1/2_ (half-time of maximal recovery) for each of the of the recovery profiles and plotted for different conditions.

EGFR diffusion coefficient was computed by averaging the normalized recovery intensity and the average was fitted with one phase association, least squares fit to compute t_1/2_

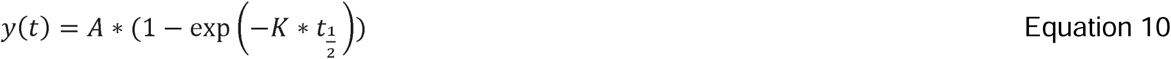

t_1/2_ was used to calculate the diffusion coefficient of EGFR-GFP on the membrane using Soumpasis equation^86,87^ and K was the rate constant.

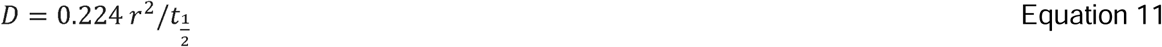

Where D is the diffusion coefficient, r is the radius of the circular FRAP area and t_1/2_ half maximal recovery time.

### Rectangular FRAP and flow analysis

A rectangular area spanning the width of the leader bleb was bleached in the rear quadrant of the bleb and intensity recovery was analyzed by drawing a line perpendicular to the long axis of the rectangle in Fiji. The intensity linescans were measured over time and each was fitted with an inverse non-linear gaussian curve fit using a Gaussian amplitude (GaussAMp) function in Origin Lab to find the minima of the fitted inverse gaussian, where the function is given by

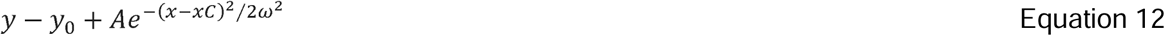

xC= center of the peak, y_0_= offset, w=width, A=amplitude of the gaussian, the position of the center of the minima of the gaussian was tracked in the images taken in the first few seconds of intensity recovery and then averaged across several blebs.

### Whole bleb FRAP and recovery analysis

An area corresponding to the whole bleb was bleached and the intensity recovery was analysed by drawing a ROI line segment across the bleb spanning from the rear to the tip of the leader bleb for recovery across the whole bleb. The measured intensity was normalized to the maximum intensity and was divided as a function of bleb length into 21 bins, taking 0 for the bleb base and 1 for bleb tip. Intensity before bleaching was taken as pre-bleach intensity and the last time point of recovery was after 100 seconds. The average from multiple such blebs were taken and ratio of last time point of recovery to the pre-bleach was plotted along with the pre-bleach and recovery intensities. For analysis of the intensity recovery rate constant, a small circular ROI of 1µm was drawn close to the neck of the bleb and the recovery intensities were recorded. The extracted intensities from each ROI was then uploaded in EasyFRAP-web FRAP analysis tool ^85^. Intensity values were normalized, and the rate constant was computed from the average by fitting with one phase association, least squares fit to compute K (as from equation 10).

#### Characteristic bleb length and width measurements

To measure the characteristic length of the bleb, a line segment was drawn from the base to the tip of the bleb in Fiji (ImageJ) and length was extracted and average of several such leader bleb lengths was taken as characteristic bleb length. For the characteristic width of the bleb, each bleb was divided into 10 equal reference points and a line segment was drawn at each point spanning across the width of the bleb and this value was extracted for the width at each of the reference point for several blebs. An average of the width across each of the reference points were reported as a characteristic width measurement.

### Actin PIV analysis

The motion of the F-tractin FusionRed-labeled actin network was determined in consecutive frames ∼6 sec apart in PIVLab ^88^ MATLAB (Mathworks). The background and the cell body were excluded from the analysis by manually masking it out. Vector velocities were used to calculate magnitudes in the leader bleb.

### Bleb elongation measurement and heat map

As the bleb protruded and elongated, the intensity measurements were made by drawing a line segment in Fiji (ImageJ) from the rear-to-front of the bleb manually. Intensity measurements were extracted and the heat map was plotted in Origin lab.

### Mathematical modelling and solutions

*Membrane protein flux and material balance:* We consider that our membrane protein (density *P*) is subject to diffusion, advection, and turnover in the bleb; based on evidence, we assume that all synthesis of the protein occurs in the cell body, and therefore there is a net transport of protein from the cell body into the bleb. The apparent diffusivity (*v_p_*), the advection velocity (*v_p_*), and turnover frequency (*k_p_*) of the protein potentially vary with position *x* in the bleb, measured from the neck of the bleb. In the material balances, we also need to account for the geometry of the bleb; its length, *L* (set to 45 μm), and position-dependent width, *w(x)* (see below), were estimated from mean measurements. Flux (*N_p_*) and steady-state material balance equations for the protein may thus be written as

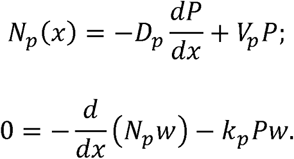

The spatial dimension *x* is defined as the distance from the neck of the bleb (μm); *x= L* is at the tip of the bleb.

*N_p_* and *v_p_* point in the direction of *x* by convention, and therefore a positive/negative value indicates net flux or flow towards the tip/neck. Zero net flux at the tip dictates the boundary condition,

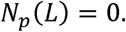

The calculated protein density profiles are presented on a normalized scale, and therefore the second boundary condition is arbitrary.

*Protein advection velocity, diffusivity, and turnover:* To further define the aforementioned *v_p_(x)* and *k_p_(x)*, it was necessary to consider the material balance of the bulk membrane. Defining *v_m_(x)* and *k_m_(x)* as the flow velocity and turnover frequency of bulk membrane, respectively, and with the ansatz that the total mass density is constant throughout the bleb (confirmed by experiment),

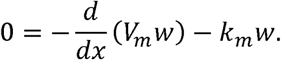

If the bleb has a constant shape, it implies that there is zero flow at the tip (*v_m_(L) = 0*). This boundary condition dictates that *v_m_ (x) c,0*; membrane flow is towards the tip of the bleb. For the calculations presented here, we took *k_m_(x)* = constant and set its value to 0.003 s^-1^, thus approximately matching the estimated bulk flow velocity of *v_m_* ≈ 0.1 μm/s measured at *x* ≈ 10 μm. Relative to the calculated *v_m_ (x)*, we consider that *v_p_(x)* is potentially influenced by two distinct, protein-corralling effects: restriction of free protein mobility, and physical interaction with the F-actin retrograde flow. Defining *v_F_ (x)* as the local velocity of the F-actin, we take

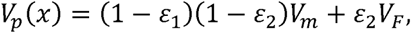

where *E_1_(x)* are *E_2_(x)* are variables, each ranging between 0 and 1, which we refer to as *corralling efficiencies*.

Their values reflect the ‘strengths’ of protein mobility restriction and interaction with flowing F-actin, respectively. The function, *v_F_* (*x*) = -0.10 + 0.08*e*^-0.2*x*^ (μm/s), was used to approximate the experimentally estimated profile from PIV analysis. Regarding the protein diffusivity, it is reduced by the corralling effects as well, and so we likewise take

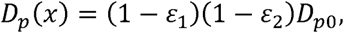

where *v_pO_* is the diffusivity value in the absence of corralling; for each scenario, the value of *v_pO_* was set to approximately match the estimated EGFR diffusivity *v_p_* ≈ 0.02 μm^2^/s measured at *x* ≈ 10 μm. Considering the potential for spatially varying *E_i_ (x)* (with *i* = 1 or 2) in a simple manner, we allow each function to be characterized by its value evaluated at the neck (*x= 0*), *E_i,O_*, and another parameter, *ε_i_*_,0_, the length scale of its gradient:

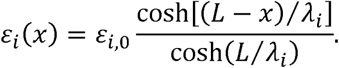

In the limit of λ_i_→ ∞, *ε_i_*(*x*) = *ε_i_*_,0_ (constant). Finally, to model protein turnover frequency in the simplest manner, consistent with observations, we equated it to that of the bulk membrane.

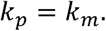

*Solution scheme:* The scheme for solving the model equations was to transform them into a set of coupled, first-order differential equations:

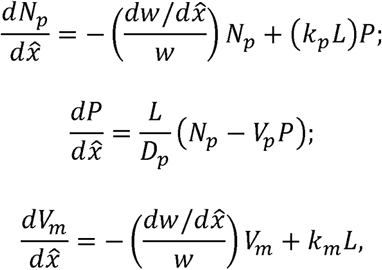

where

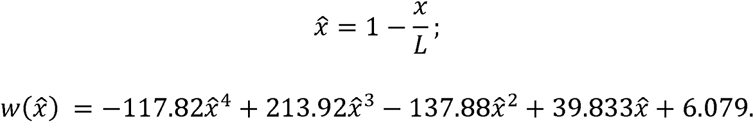

The latter equation, was fit to the mean of 17 representative width profiles (μm). This scheme permitted forward integration, with boundary conditions

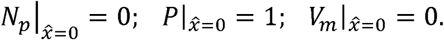

The domain was discretized with a spatial step size of *0.001*, and the initial value problem was solved using the improved Euler method/Heun’s method, implemented in Excel for rapid evaluation of the model output and analysis of numerical accuracy and stability. Once the solution for each scenario was obtained, the *1(x)* profile was normalized by the maximum value for presentation in figures. Figure 5a, EGFR density (P) with changes in *E_1_(x)* which reflect the ‘strengths’ of protein mobility restriction. It was set to 0.1, 0.3, 0.5, 0.7 and 0.9 while *E_2_(x)* which reflect the interaction with flowing F-actin was set constant to 0.1 (weak interaction). The actin retrograde flow velocity was set to *v_F_* (*x*) = -0.10 + 0.08*e*^-0.2*x*^ (μm/s).

Extended Figure 2l, EGFR density (P) with changes in actin retrograde flow where measured *v_F_* (*x*) = -0.10 + 0.08*e*^-0.2*x*^ (μm/s) was taken optimum. Following retrograde flow velocities were considered: 0.01, 0.05, 0.1 (optimum), 0.15, 0.2 (μm/s). *E_1_(x)* which reflect the ‘strengths’ of protein mobility restriction was set to 0.1 (weak) while *E_2_(x)* which reflect the interaction with flowing F-actin was set constant to 0.5 (strong)

### Statistical analysis

Statistical analysis was performed in Prism (GraphPad). Graphs represent mean values ± SD as indicated in the figure legends of atleast three independent experiments. Statistical significance was obtained using student t-test with Welch’s, Kolmogorov-Smirnov or Mann-Whitney correction as indicated in the figure legends.

Significance is reported in the figure legends with P>0.05 being n.s.

## Supporting information

Modelling supplemental text

## Scientific illustrations

Illustrations and cartoons were made using Adobe illustrator and BioRender.

## Acknowledgements and funding

We thank the Waterman Lab especially William Shin, Dr. Ana Pasapera, the National Heart, Lung, and Blood Institute (NHLBI), the NHLBI light microscopy (Xufeng) and flow cytometry core facilities. We would also like to thank Dr. Tamas Balla, Dr. Senthil Arumugam, Dr. Valentine Jaumouille for their feedbacks and discussions on this work. This work was funded by the NHLBI Division of Intramural Research (DIR) (AJ and CMW), National Institute of General Medical Sciences under award number R01 GM141691 (JMH), the National Institute of Biomedical Imaging and Bioengineering of the National Institutes of Health under award number U01 EB018816 (JMH), NIH grant 5R01GM144362 and a Vallee Scholar award (JET).

---

**Extended Figure 1:**
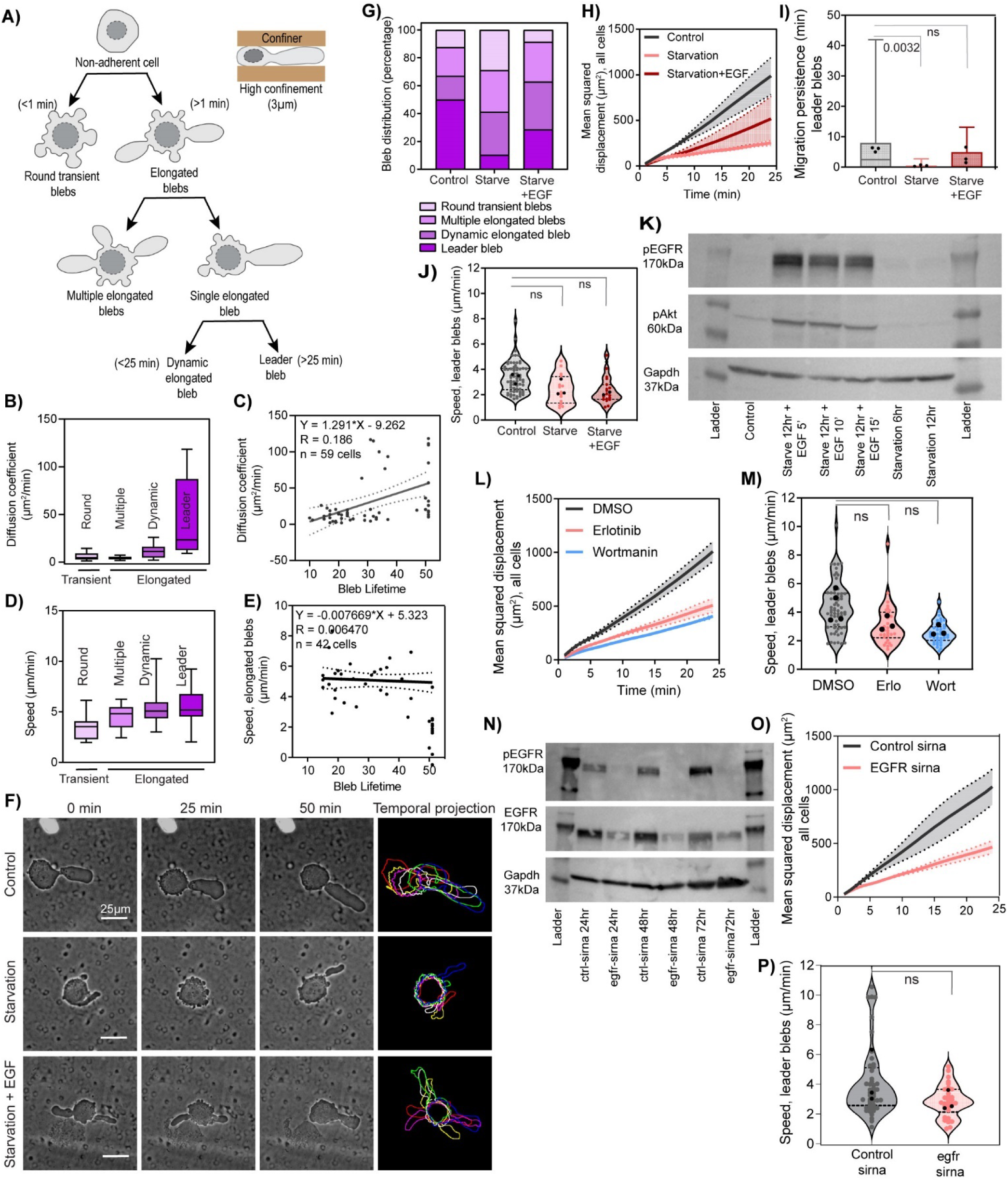
Leader bleb-based migration requires EGFR signaling. (A) Schematic diagram illustrating the cell confinement geometry used in this study (top right) and the classification scheme of different bleb morphologies seen when A375M melanoma cells are subjected to high confinement and low adhesion (left). Under such confinement, cells exhibit either round transient blebs or elongated blebs. Based on the number of elongated blebs, cells were sub-classified into multiple or single elongated blebs. Cells with single elongated blebs were further sub-classified based on the single elongated bleb lifetime into dynamic (<25 min) and leader bleb (>25 min). (B-E) Analysis of confined cells in the bleb morphology classes defined in (A). Round transient blebs (n= 21 cells), multiple elongated blebs (n= 18 cells), dynamic elongated blebs (n= 31 cells), and leader blebs (n= 44 cells) from N=3 experiments. (B) Box plots of diffusion coefficient. (C) Diffusion coefficient of cells exhibiting polarized elongated blebs relative to bleb lifetime (n=59 cells with elongated blebs). (D) Box plot of speed of different bleb phenotypes in high confinement. (E) Average speed of cells exhibiting a single elongated bleb compared to the bleb lifetime (n=55 elongated blebs), solid black line, linear fit (F) Representative time-lapse phase-contrast images (left three columns) of A375M2 melanoma cells under 3 µm confinement, temporal color-encoded projection of cell shapes at 10-minute intervals (right column) of the cells shown at left. Cells were either untreated (control, top row) serum starved (starvation, middle row) or serum starved then stimulated with EGF (starvation + EGF, bottom row) addition compared to the controls with FBS (left panel). (G-J) Analysis of confined cells after starvation and EGF addition (Control, n=112 cells, Starvation, n=97cells and Starvation+EGF, n=70 cells). (G) Quantification of bleb phenotype distribution percentage. (H) Mean-square displacement over time with average (dark line) and standard deviation (dotted line, shaded area) shown. (I) Box plot of migration persistence of cells with leader blebs (Control, n=56 cells, Starvation, n=14 cells and Starvation+EGF, n=18 cells). (J) Violin plot of migration speed. (K, N) Western blots of lysates of A375M cells, molecular weight noted in kDa. (K) Cells were either untreated (Control) or starved for 6 or 12 hours. 12 hour starved cells were restimulated with EGF for 5, 10 and 15 minutes. Blots probed for pEGFR, pAkt and Gapdh as a loading control. (L, M) Analysis of confined cells treated with vehicle (DMSO), erlotinib (50nM) or wortmannin (10μM). (DMSO, n=64 cells, erlotinib, n=30 cells, wortmannin, n=39 cells) (L) Mean-square displacement over time with average (dark line) and standard deviation (dotted line, shaded area) shown. (M) Average migration speed of cells undergoing leader bleb-based migration. (N) Cells were either untreated transfected with non-targetting control siRNAs (ctrl-sirna) or siRNAs targeting EGFR (egfr sirna) for 24, 28, or 72 hrs. Blots probed for pEGFR, total EGFR, and Gapdh as a loading control. (O, P) Analysis of confined cells after transfection with non-targetting siRNA or siRNA targeting EGFR (control siRNA n=48 cells; EGFR siRNA n=30 cells). (O) Mean-square displacement over time with average (dark line) and standard deviation (dotted line, shaded area) shown. (P) Average migration speed of cells undergoing leader bleb-based migration. Scale bars= 25 microns.

**Extended Figure 2:**
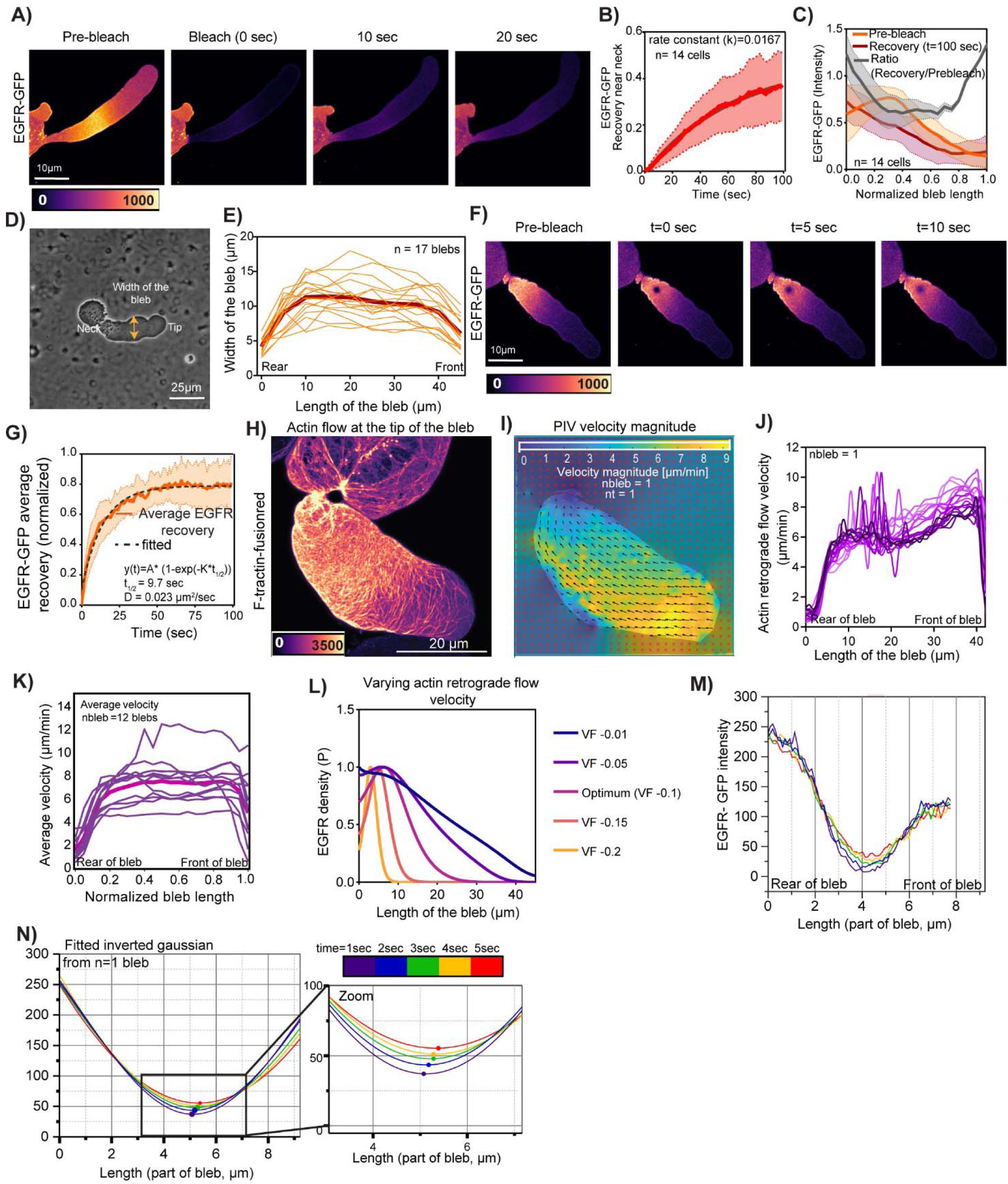
EGFR and other transmembrane proteins form rear-to-front gradients on leader blebs. (A, F, H) Confocal and (D) phase-contrast images of A375M2 melanoma cells under 3μm confinement. A) Pseudocolored intensity image of cells expressing EGFR-GFP that were subjected to photo-bleaching of thenleader bleb (left-center panel) and time-lapse imaging during fluorescence recovery (right 2 panels). (B). Average normalized fluorescence intensity over time (solid lines) +/- standard deviation (dotted lines and shaded area) from small areas near the bleb neck in photobleached regions of cells subjected to photobleaching like that shown in (A), Rate of recovery k=0.0167. (C) Mean normalized intensity profile (dark lines) of EGFR-GFP across the normalized length of the leader bleb, ± s.d. (dotted lines, shaded areas), from linescan analysis of cells under the treatment shown in (A). Time zero is the first image after photobleaching. Ratio of recovery intensity to the pre-bleach intensity is plotted in the same graph (n=14 cells). (D) Cell undergoing leader bleb based migration under high confinement illustrating parameters plotted in (E) Bleb width as a function of position along the bleb, light orange= individual cells, dark orange= average, (n=14 cells). (F) Pseudocolored intensity image of cells expressing EGFR-GFP that were subjected to photo-bleaching of small area near the base of the bleb (t= 0 sec) and time-lapse imaging during fluorescence recovery. (G) Average normalized fluorescence intensity over time (solid line) from photobleached regions like that shown in (F), time zero is the first image after photobleaching. Single exponential least squares fits (dotted lines). (H) Pseudocolored intensity super resolution image of a leader bleb in a cell expressing F-tractin-fusion red to label actin filaments. (I) PIV analysis of time-lapse movies of F-tractin-fusionred of the blebs shown at left showing the direction (arrows) and magnitude (color encoded) of actin retrograde flow in two consecutive frames of the movie. (J) Actin retrograde flow velocities as a function of position along the bleb from a single bleb over time (light purple, t=0sec to dark purple, t=120 sec) (K) Actin retrograde flow velocities as a function of normalized position along the leader bleb. Purple lines= individual cells, magenta line= average of n=12 cells. (K) Outcome of the model. EGFR density with varying actin retrograde flow velocities, with optimum being the measured actin retrograde flow velocity. (M) Intensity line-scans from the highlighted position in figure (4) taken at one-second intervals (color encoded). (N) Left and Right: Each graph (from M) was fitted with an inverse gaussian and the minima of the fit is marked. Right: Zoom of the minima region of the graph.

**Extended Figure 3:**
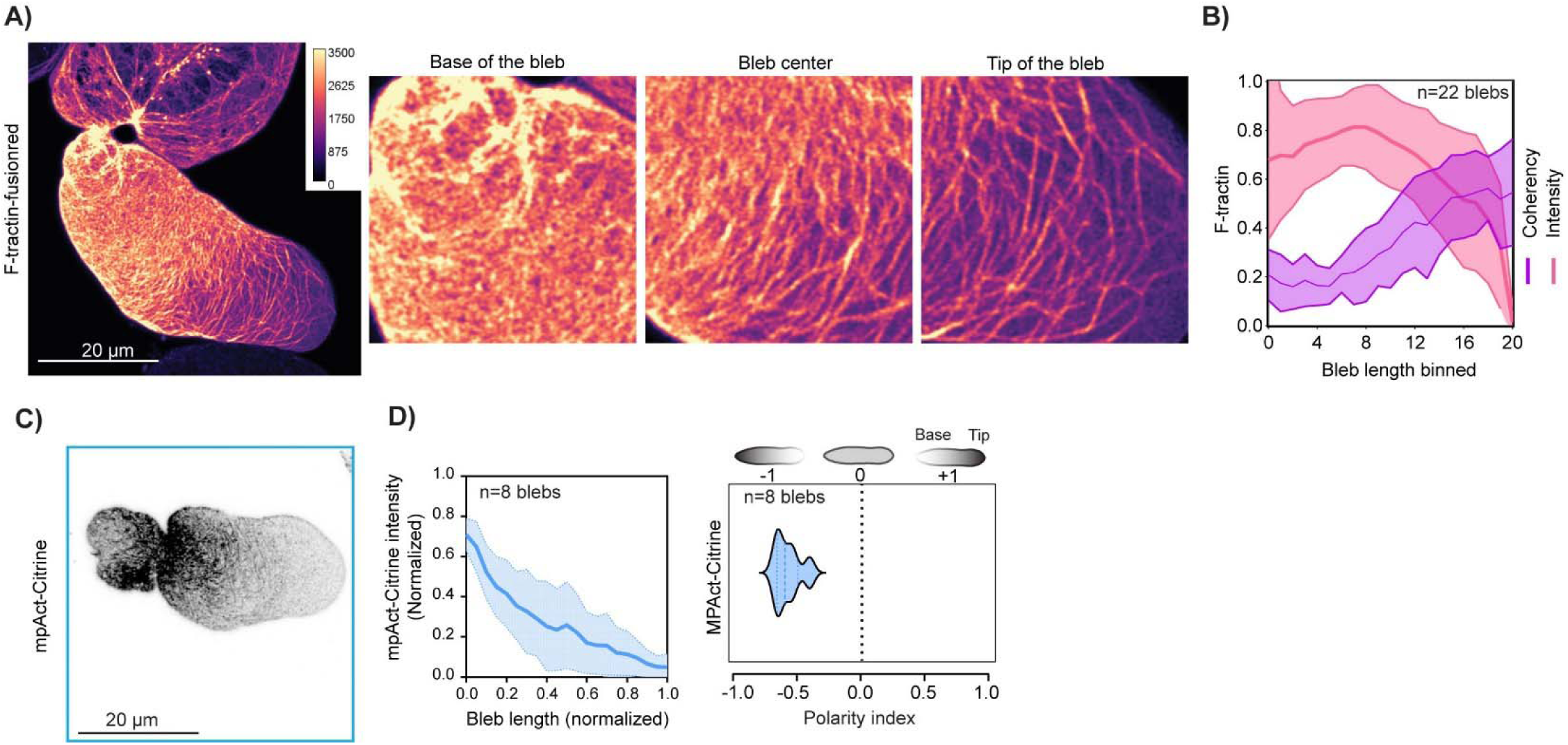
A dense actin meshwork forms close to the plasma membrane at the rear of leader blebs. (A, C) Confocal images of A375M2 melanoma cells under 3μm confinement. (A) Left: Pseudocolored intensity super resolution image of cell expressing F-tractin-fusionRed. Right panels: Zooms of the noted regions of bleb in the cell at (right), showing the actin architecture at the base (left center), center (right center) and tip (right) of the bleb. (B) Average normalized intensity (red) and actin filament coherency (purple) (dark lines) +/- standard deviation (light lines and shaded areas) or F-tractin fusionred as a function of binned position along the bleb length, n=22 cells. (C) Inverted grayscale image of mpAct-Citrine. (D) Left panel: Average normalized intensity (dark line) +/- standard deviation (light lines and shaded areas) of cells expressing mpAct-citrine like in (C) as a function of position along the normalized bleb length, right panel: Violin plot of polarity index of mpAct-citrine in blebs. N=8 cells from 3 experiments.

**Extended Figure 4:**
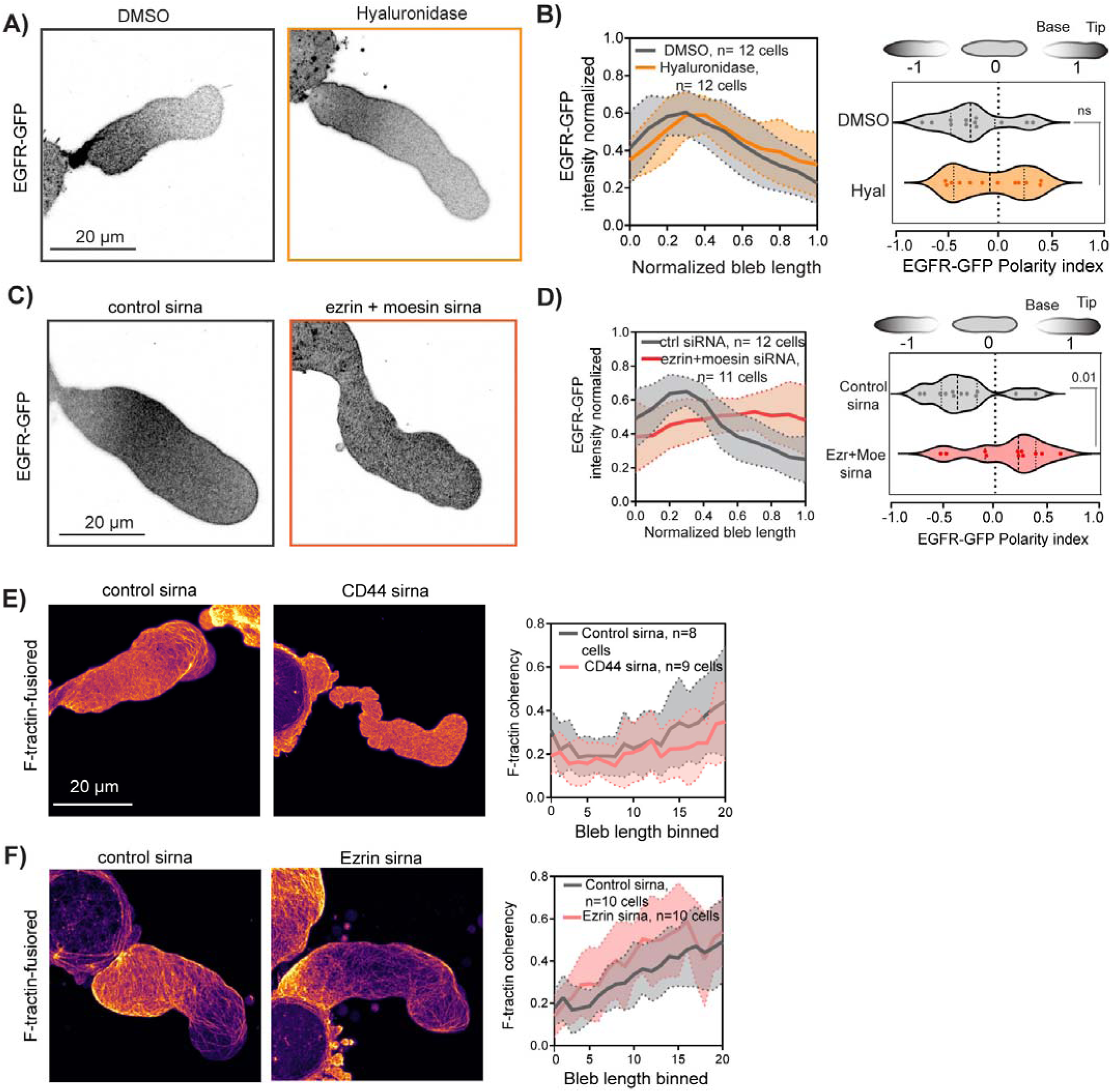
CD44 corralling is independent of its receptor activity. (A, C, E, F) Confocal images of A375M2 melanoma cells under 3μm confinement. (A) Inverted contrast image of cells expressing EGFR-GFP treated with either DMSO or Hyaluronidase. (B) Left panel: Average normalized intensity (dark lines) +/- standard deviation (light lines and shaded areas) of EGFR-GFP as a function of normalized position along the normalized bleb length in cells treated with DMSO (gray, n=12 cells) or Hyaluronidase across (orange, n=12 cells). Right panel: Violin plot of polarity indices of EGFR-GFP in blebs of the cells analyzed at left, Welch’s t-test p=n.s. (C) Inverted contrast image of cells expressing EGFR-GFP co-transfected with control-siRNA or ezrin and moesin-siRNA. (D) Average normalized intensity (dark lines) +/- standard deviation (light lines and shaded areas) of EGFR-GFP as a function of normalized position along the normalized bleb length in cells co-transfected with control-siRNA (gray, n=12 cells) or ezrin and moesin-siRNA (red, n=11 cells). Right panel: Violin plot of polarity indices of EGFR-GFP in blebs of the cells analyzed at left, Welch’s t-test p=n.s (E) Left and Center panels: Pseudocolored intensity image of cells expressing F-tractin-FusionRed and co-transfected with control-siRNA (left) or CD44-siRNA (center). Right: Average normalized F-actin coherency (dark lines) +/- standard deviation (light lines and shaded areas) of F-tractin localization as a function of normalized position along the normalized bleb length in cells co-transfected with control-siRNA (gray, n=8 cells) or CD44-siRNA (pink, n=9 cells). (F) Left and Center panels: Pseudocolored intensity image of cells expressing F-tractin-FusionRed and co-transfected with control-siRNA (left) or ezrin-siRNA (center). Right: Average normalized F-actin coherency (dark lines) +/- standard deviation (light lines and shaded areas) of F-tractin staining as a function of normalized position along the normalized bleb length in cells co-transfected with control-siRNA (gray, n=10 cells) or ezrin-siRNA (pink, n=10 cells).

**Extended Figure 5:**
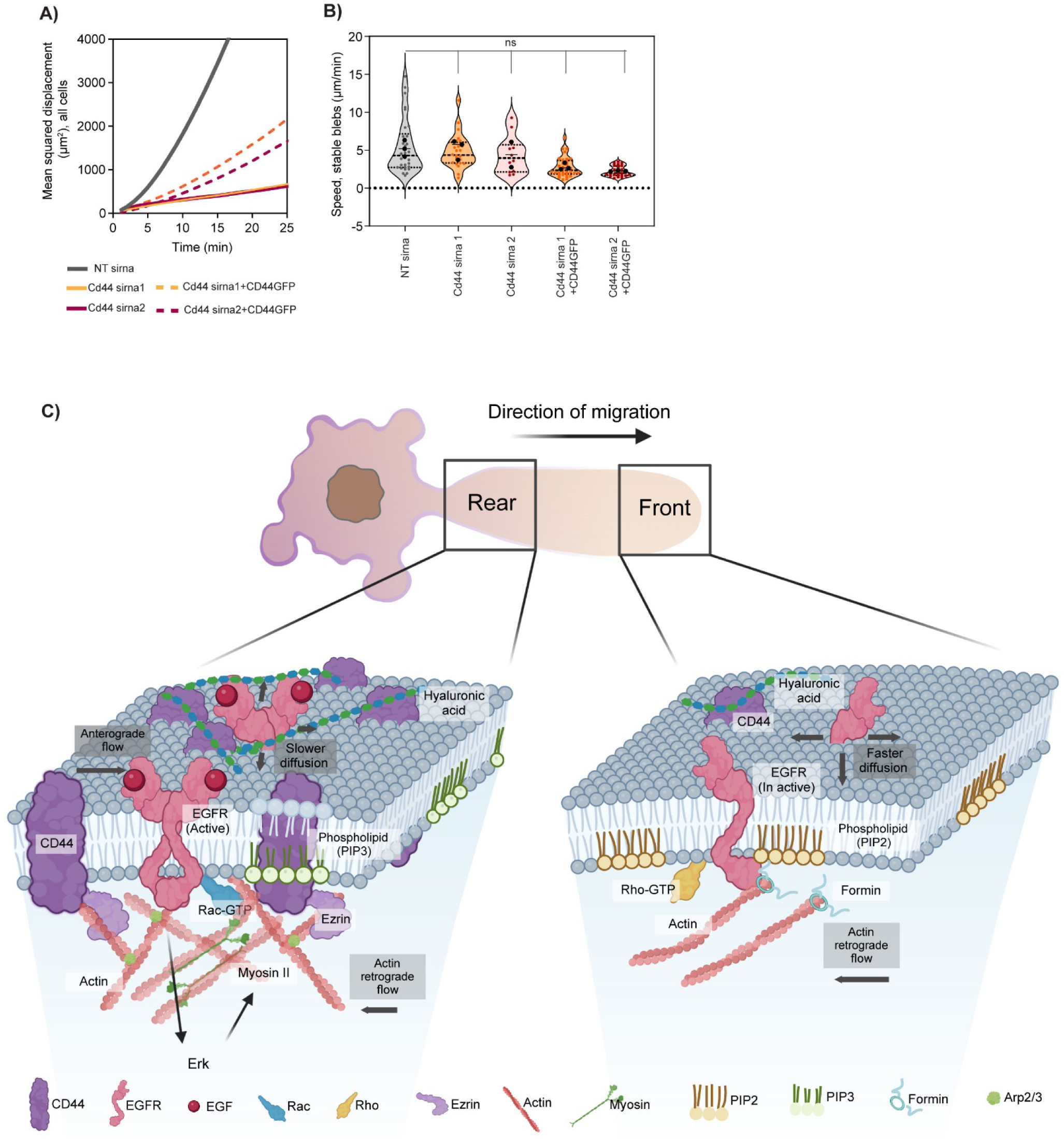
Model of molecular regulation of EGFR polarity during leader bleb-based migration under non-adhesive confinement. (A, B) Quantitative analysis of cell migration tracks from time-lapse phase contrast movies of A375M cells under non-adhesive confinement to 3 Lm like those shown in (Fig 7C). (B) Average mean-square displacement over time. (Control-sirna, gray solid line, n=47 cells, CD44-siRNA1, solid orange line, n=122 cells, CD44-siRNA2, dotted orange line, n=69 cells, CD44-siRNA1 + CD44-GFP, solid maroon line, n=121 cells, CD44 siRNA2 + CD44-GFP, dotted maroon line n=75 cells). (B) Violin plots of average migration speed. Control-siRNA, gray, n=25 cells; CD44-siRNA1, orange dots, n=10 cells; CD44 siRNA2, maroon dots, n=12 cells; CD44-siRNA1 + CD44-GFP, orange boxes, n=20 cells; CD44siRNA2 + CD44-GFP maroon boxes, n=17 cells. (C) Hypothetical model showing molecular mechanism of maintenance rear-tor front gradient of EGFR activity during leader bleb based migration. See discussion.

