## Supplementary material for "CD44 and Ezrin restrict EGF receptor mobility to generate a novel spatial arrangement of cytoskeletal signaling modules driving bleb-based migration": Modelling supplemental text

**Supplementary text**

**Mathematical model formulation**

**1. Initial formulation of the model equations from general conservation principles**

For the membrane protein of interest, with local density $P(x)$, its transport is characterized by local flux in the $x$ direction, $N_{p}\left( x \right)$, defined according Fick’s law of diffusion plus the potential contribution of advective flow:

$$N_{p}\left( x \right)=-D_{p}\frac{dP}{dx}+V_{p}P.$$

In this equation, $D_{p}(x)$ is the local protein diffusivity, and $V_{p}(x)$ is the local advection velocity of the protein; for the sake of generality, both are allowed to vary with position $x$ at this point. Considering the geometry of the bleb, defined by length, $L$, and width, $w(x)$, the steady-state material balance for the membrane protein is

$$0=-\frac{d}{dx}\left( N_{p}w \right)+R_{p}w,$$

where $R_{p}(x)$ is generally defined as the local rate of protein generation (net insertion into the membrane less the loss from the membrane, regardless of the mechanism(s) involved). A similar steady-state balance is invoked for the bulk membrane (mass density $M$), except for the consideration that there is, by definition, no diffusion of total mass:

$$0=-\frac{d}{dx}\left( V_{m}Mw \right)+R_{m}w.$$

In this equation, $V_{m}(x)$ is the local bulk membrane velocity, and $R_{m}(x)$ is the local rate of bulk membrane generation (net addition minus loss). Based on this formulation, the quantities ${-R}_{p}/P$ and $-R_{m}/M$ are defined as the local turnover frequencies of the membrane protein and of the bulk membrane, respectively.

**2. Derivation of conditions for maintenance of any protein gradient**

Starting from the general formulation posed above, we stipulate (supported by experimental observations, that the mass density of the bulk membrane is uniform ($M$ is constant). Hence, we manipulate the balance on bulk membrane to obtain,

$$0=-\frac{d}{dx}\left( V_{m}w \right)+\frac{R_{m}}{M}w,$$

and again by simply multiplying both sides by $P(x)$:

$$0=\left( -\frac{d}{dx}\left( V_{m}w \right)+\frac{R_{m}}{M}w \right)P.$$

Turning to the protein balance, we incorporate the flux expression to obtain,

$$0=-\frac{d}{dx}\left( N_{p}w \right)+R_{p}w=\frac{d}{dx}\left( D_{p}w\frac{dP}{dx} \right)-\frac{d}{dx}\left( V_{p}wP \right)+R_{p}w.$$

Manipulating the above, to separate the terms that contain the protein gradient, $\frac{dP}{dx}$,

$$\frac{d}{dx}\left( D_{p}w\frac{dP}{dx} \right)-V_{p}w\frac{dP}{dx}=P\frac{d}{dx}\left( V_{p}w \right)-R_{p}w.$$

Next, we add the bulk membrane balance (which is equal to zero on both sides) to the above:

$$\frac{d}{dx}\left( D_{p}w\frac{dP}{dx} \right)-V_{p}w\frac{dP}{dx}=P\frac{d}{dx}\left( V_{p}w \right)-R_{p}w+\left( -\frac{d}{dx}\left( V_{m}w \right)+\frac{R_{m}}{M}w \right)P.$$

Combining the terms on the right-hand side, we obtain,

$$\frac{d}{dx}\left( D_{p}w\frac{dP}{dx} \right)-V_{p}w\frac{dP}{dx}=\left\{ \frac{d}{dx}\left[ \left( V_{p}-V_{m} \right)w \right]-\left( \frac{R_{p}}{P}-\frac{R_{m}}{M} \right)w \right\}P.$$

Inspection of the above indicates that, if there is no gradient ($\frac{dP}{dx}=0)$, the right-hand side of the equation must be equal to zero. For this to be true throughout the domain, the only plausible scenario is

$$V_{p}\left( x \right)=V_{m}\left( x \right); \text{and, }\frac{R_{p}(x)}{P}=\frac{R_{m}(x)}{M}.$$

This scenario may be stated thus: the membrane protein both flows with, and is turned over at the same frequency as, the bulk membrane. If this were true, then the proof of zero gradient throughout can be formalized; transforming the resulting equation into a first-order differential equation yields,

$$\frac{dY}{dx}-\frac{V_{p}}{D_{p}}Y=0; Y=D_{p}w\frac{dP}{dx}.$$

Given the boundary conditions,

$$N_{p}\left( L \right)=0; V_{p}\left( L \right)=V_{m}\left( L \right)=0,$$

we conclude that $Y\left( L \right)=w\left( L \right)\left[ -N_{p}\left( L \right)+V_{p}\left( L \right)P\left( L \right) \right]=0$, and therefore the trivial solution is $Y\left( x \right)=0$.

For the scenarios considered in this paper, we invoked the basic assumption,

$$\frac{R_{p}(x)}{P(x)}=-k_{p}=\frac{R_{m}\left( x \right)}{M}=-k_{m}.$$

In words, membrane protein (EGFR) is removed from the bleb at the same (constant) frequency as (along with) bulk membrane. A difference between the advective flow velocities of EGFR and bulk membrane ($V_{p}\left( x \right)\neq V_{m}\left( x \right)$) is sufficient to predict a gradient of EGFR in the bleb.
